# Paralog interference contributes to the preservation of genetic redundancy

**DOI:** 10.1101/2025.06.13.659495

**Authors:** Angel F. Cisneros, Florian Mattenberger, Isabelle Gagnon-Arsenault, Yacine Seffal, Lou Nielly-Thibault, Alexandre K. Dubé, María del Mar Varela Vásquez, María Camila Muñoz Vega, Christian R. Landry

**Affiliations:** Département de biochimie, de microbiologie et de bio-informatique, Faculté des sciences et de génie, Université Laval, G1V 0A6, Québec, Canada; Institut de biologie intégrative et des systèmes, Université Laval, G1V 0A6, Québec, Canada; PROTEO, Le regroupement québécois de recherche sur la fonction, l’ingénierie et les applications des protéines, Université Laval, G1V 0A6, Québec, Canada; Centre de recherche sur les données massives, Université Laval, G1V 0A6, Québec, Canada; Gregor Mendel Institute of Molecular Biology, Austrian Academy of Sciences, 1030, Vienna, Austria; Département de biologie, Faculté des sciences et de génie, Université Laval, G1V 0A6, Québec, Canada; Departamento de Ciencias Biológicas, Bioprocesos y Biotecnología, Facultad Barberi de Ingeniería, Diseño y Ciencias Aplicadas, Universidad Icesi, 760008, Cali, Colombia; Departamento de química, Facultad de ciencias naturales y exactas, Universidad del Valle, 760032, Cali, Colombia

**Keywords:** Gene duplication, loss of function, protein complexes, paralog interference

## Abstract

Models of gene duplication often assume that loss-of-function mutations neutrally promote the return to the ancestral singleton state. They thus ignore the potential functional interference between duplicated proteins stemming from their physical interactions. Here, we show that for heteromerizing paralogs, such interference potentiates negative selection on loss-of-function mutations. This effect maintains genetic redundancy over longer timescales depending on the rate and severity of loss-of-function mutations. We experimentally estimate that around 6% of deleterious amino acid substitutions for a representative tetrameric protein interfere with a second copy. Interfering substitutions typically disrupt either catalysis or the final step of protein complex assembly, with varying degrees of severity. Our work shows that paralog interference renders the negative effects of loss-of-function mutations visible to purifying selection, contributing to the preservation of genetic redundancy.

## Introduction

Gene duplication is the main source of genetic redundancy^1–3^. Immediately after a duplication event, the two paralogs are usually functionally redundant as they have identical sequences and regulatory elements. A common assumption is that loss-of-function (LOF) mutations in one copy are rendered neutral by the presence of its paralog. Since LOF mutations occur much more frequently than other types of mutations^4,5^, the most likely long-term fate of duplicated genes is thought to be a return to a single functional copy. Alternative fates include the divergence of functions between paralogs leading to new functions^6^ or to the partitioning of ancestral ones^7^.

These two scenarios happen gradually and require the persistence of the two copies in the genome for long enough so rare function-affecting mutations could occur. Therefore, the timescales of maintenance of redundant duplicates have consequences for the robustness of cells to mutations and for adaptive evolution. More generally, the number of genes in the genome depends on the balance between the rates of duplication and gene loss^8^. Any biochemical or biophysical factor that decreases the rate of gene loss will lead to the maintenance of a larger number of genes and the expansion of gene families in the long term.

Previous studies have identified factors that affect the rate of gene loss following gene duplications. For example, genes encoding self-interacting (homomeric) proteins^9,10^ and those more likely to acquire dominant deleterious mutations^11,12^ appear to be preferentially retained. These two observations could be mechanistically connected, since the duplication of genes encoding homomeric proteins introduces paralogous proteins that physically interact with each other^13,14^. Classical models for the retention of gene duplicates assume that LOF impacts only the gene copy bearing a mutation, i.e. that the genes contribute independently to function, as in the case of duplicates interacting with a common interaction partner^15^. However, because of their physical interactions, duplicated homomeric proteins are not independent. As a result, LOF mutations in one paralogous protein could interfere with the normal function of the other one, causing a stronger decrease in function than LOF mutations would cause for non-interacting paralogs (Fig. 1A-C). Here, we show that negative selection against such LOF mutations could preserve the two interfering paralogs in the initial redundant state for a longer period than paralogs that do not interact (Fig. 1B-C). Since this phenomenon would be analogous to dominant negative alleles at the same locus in diploids, whereby LOF mutations in one allele can inactivate the protein produced by the other allele due to their physical assembly^16,17^, we refer to it as negative paralog interference. Considering that at least 30% of genes code for homomeric proteins^18,19^, negative paralog interference could have a profound effect on shaping extant genomes. We use models and experiments to test if and how homomerization provides access to deleterious mutations that ultimately preserve redundancy.

**Figure 1.**
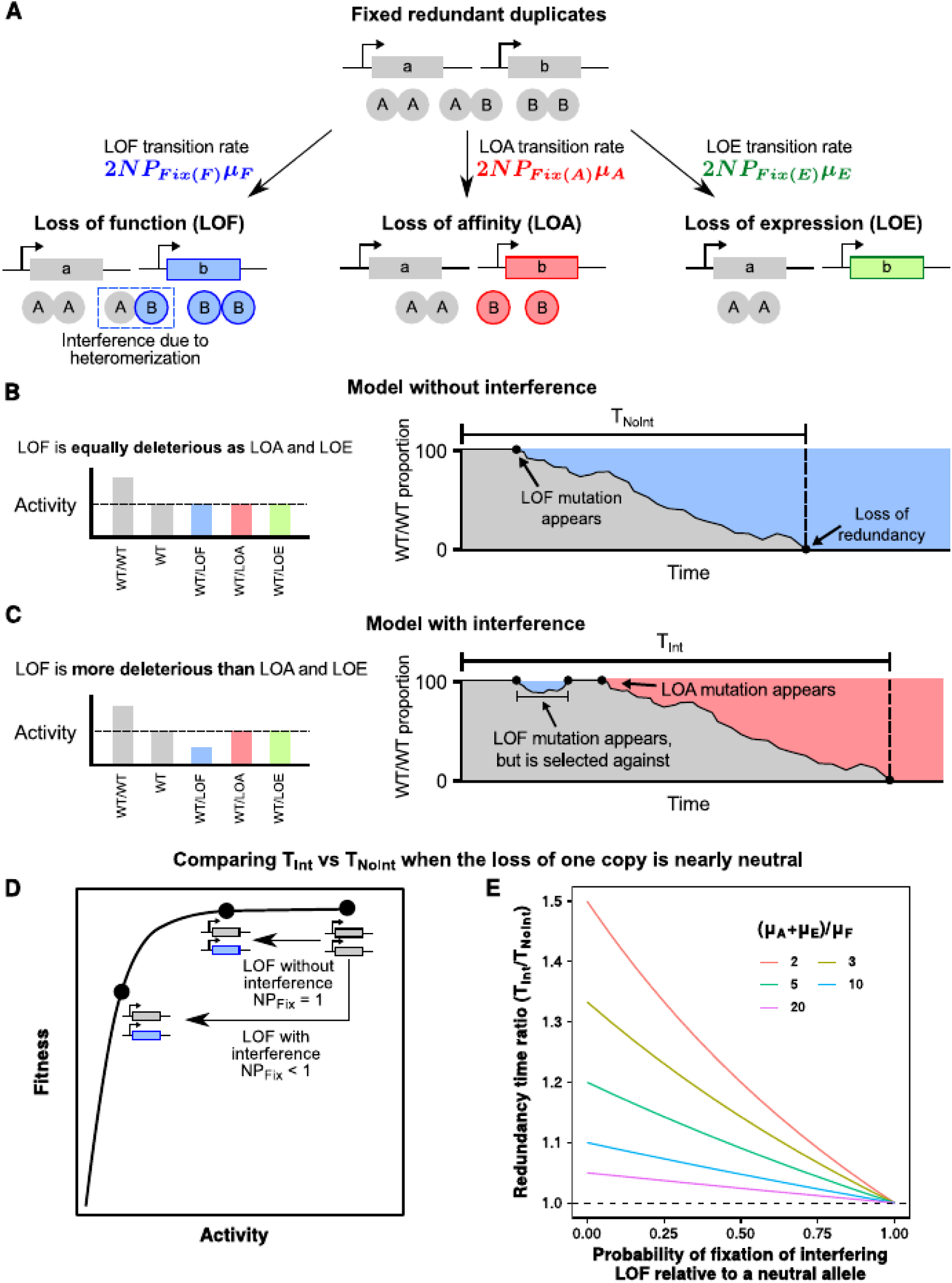
Interference between paralogs leads to longer periods of genetic redundancy. **A)** Model of the transitions from the initial state to loss of function, affinity, or expression of one copy. These transitions depend on their respective mutation rates (*µ*) and probabilities of fixation (*P_Fix_*). **B-C)** Two scenarios for the effect of interference on the maintenance of redundant paralogs. In the classical case without interference (B), LOF mutations are equally deleterious as LOA and LOE mutations. Redundancy can be lost over time as an LOF mutation drifts to fixation. With interference (C), LOF mutations are more deleterious than LOA and LOE. Thus, LOF mutations would be more strongly selected against and less likely to fix, although redundancy could be still lost if an LOA or LOE mutation fixes. **D)** Fitness function with diminishing fitness returns for increases in activity. A reduction of 50% in activity (LOE, LOA, and LOF mutations without interference) is nearly neutral, for instance if duplication was neutral in the first place, while stronger decreases (LOF mutations with interference) are deleterious. **E)** Ratio of the time needed to lose redundancy with interference (*T_Int_*) versus without interference (*T_NoInt_*). Different curves indicate varying mutation rates.

### Negative paralog interference increases the time required to lose genetic redundancy

To quantify the impact of interference, we examine transitions of populations from an initial state having two fixed redundant functional paralogs to a single functional one, that is, the loss of redundancy. We focus on genes encoding obligate homodimeric enzymes, such that three groups of mutations could eliminate their contribution to total activity and thus cellular fitness: loss-of-function (LOF), loss-of-affinity (LOA), and loss-of-expression (LOE) (Fig. 1A). These three types of mutations would respectively disrupt the catalysis, dimerization, and synthesis of the protein. Other mechanisms such as loss-of-translation or loss-of-stability could be considered but their effect on function would be the same as LOE. The transitions from a state with redundant paralogs to one where one paralog is inactivated by LOF, LOA, or LOE mutations depend on the respective mutation rates (*µ_F_*, *µ_A_*, *µ_E_*), the effective population size (*N*), and the probabilities of fixation (*P_Fix_*). Since these transitions result in the loss of redundancy, the expected time needed to lose redundancy (*T*) is:

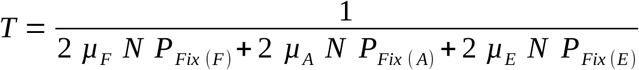

The above expression allows us to compare the times needed to lose redundancy with (*T_Int_*) and without interference (*T_NoInt_*). To do so, we calculate the ratio *T_Int_*/*T_NoInt_* using the same mutation rates and effective population size, which for simplicity we assume to be identical to the census population size (see Methods). The only parameter that changes depending on the presence or absence of interference is the probability of fixation of LOF mutations (*P_Fix(F)_*). When there is no interference, mutations causing LOF, LOA, or LOE only inactivate their respective allele, halving the total biochemical activity produced by the paralogs. Thus, they would have the same fitness effect and probability of fixation as the loss of one copy (*P_Fix(F)_ = P_Fix(A)_ = P_Fix(E)_ = P_Fix(Loss)_*). Conversely, in the presence of interference, LOF mutations could affect the function of the other allele, while LOA and LOE mutations would not be expected to, at least in the simple case of a dimeric protein that assembles in one step. Since LOA alleles cannot form any homo-or heterodimers and all the WT copies still assemble into homodimers, this scenario is similar to LOE. Accordingly, the probability of fixation of LOF mutations would be reduced as they become more deleterious on average (*P_Fix(F)_ < P_Fix(A)_ = P_Fix(E)_ = P_Fix(Loss)_*).

Typical fitness functions provide diminishing returns from total protein activity^20–22^. Thus, we first explored the scenario in which the loss of one copy is nearly neutral, that is, halving total activity does not result in a decrease in fitness (*NP_Fix(Loss)_* = 1) (Fig. 1D). In this case, the ratio *T_Int_*/*T_NoInt_*can be simplified to the following expression (see Methods):

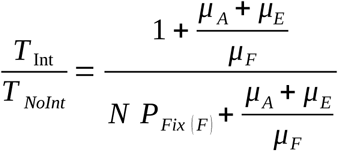

The ratio *T_Int_*/*T_NoInt_* becomes a function of the probability of fixation of the LOF allele with interference (*P_Fix(F)_*) and the ratio of mutation rates (*µ_A_* + *µ_E_*) / *µ_F_*. The ratio *T_Int_*/*T_NoInt_* increases with both decreasing probabilities of fixation of the LOF allele and decreasing ratio of mutation rates (Fig. 1E). Importantly, the expected increase in time needed to lose redundancy is accompanied by an increase in the variance, which could further potentiate the maintenance of redundant paralogs in some populations (see Methods). We also consider the case in which the loss of one copy is deleterious (Fig. S1A), and we observe an increase in the time needed to lose redundancy with decreasing values of *P_Fix(F)_*, as long as LOF mutations are more deleterious with interference than the loss of one copy (Fig. S1B, Methods). Thus, the rate and severity of LOF mutations that interfere negatively with the sister copy contribute to the preservation of genetic redundancy.

### A study system to identify and quantify the impact of interfering mutations

The rate of mutations causing negative paralog interference and their fitness consequences appear to be important determinants of excess retention time. We therefore set out to examine these parameters experimentally.

While we have focused on dimers so far, paralogous proteins can also assemble into larger oligomeric complexes. Dimeric paralogs with the same abundance and binding affinity will form a mixture of 25% of each homodimer and 50% of the heterodimer. If interfering mutations inactivate the mutant homodimers and the heterodimer, the fraction of functional complexes would be reduced to 25%. Such interfering mutations would be more deleterious than the loss of one gene copy, which decreases activity to 50%. The effects of negative interference would be even more severe for higher order assemblies. For tetramers, the expected fraction of oligomers containing only subunits without the interfering mutation is around 6% (Fig. 2A)^23^. In addition, for larger oligomeric complexes, the presence of multiple assembly interfaces could potentially make LOA also contribute to interference as partially assembled complexes could be inactive^17^.

**Figure 2:**
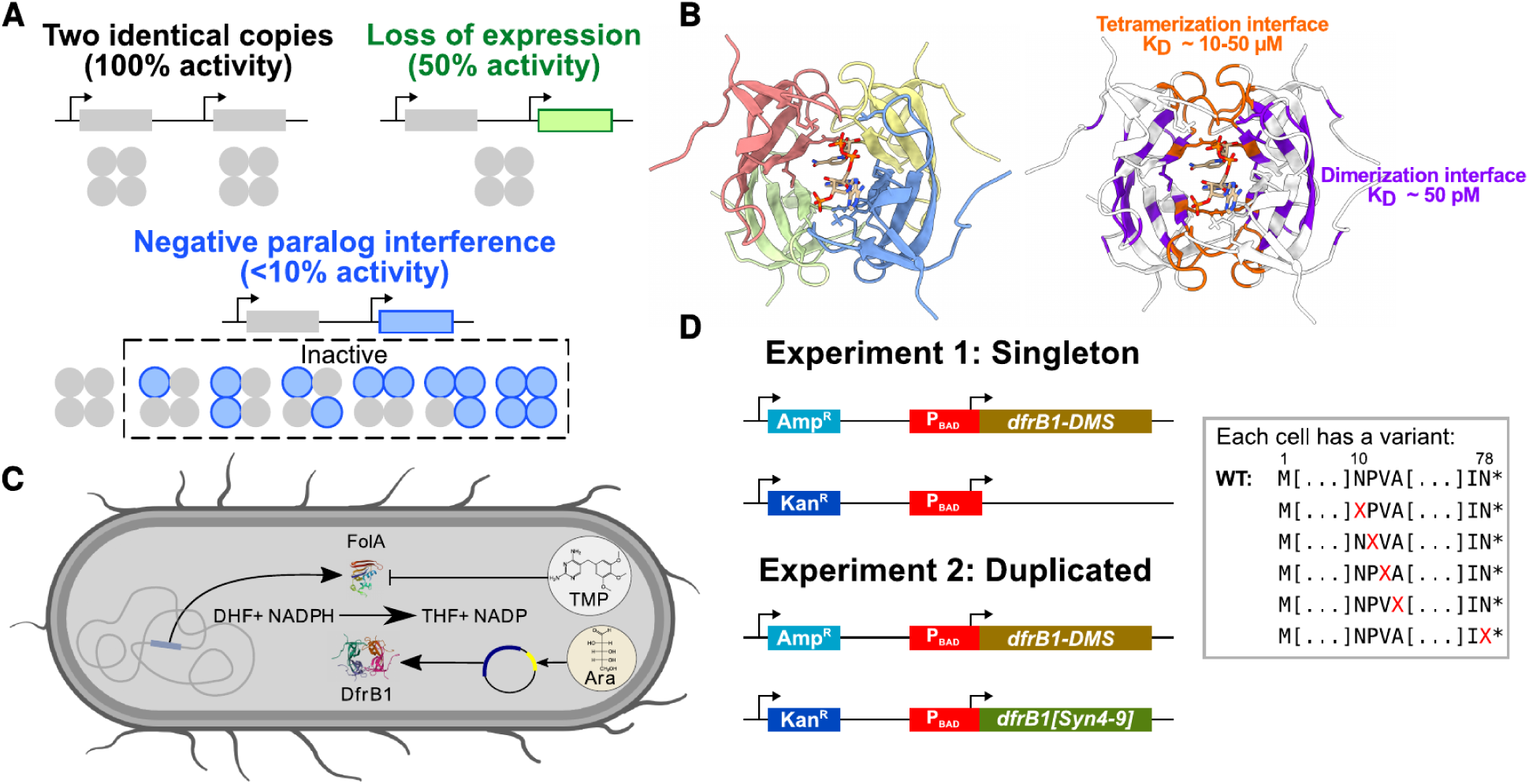
Study model to identify and measure the impact of negative interference mutations. **A)** The loss of expression of one copy halves the total concentration of complexes, while negative paralog interference produces a larger fraction (>90%) of inactive tetramers. **B)** Structure of DfrB1. *Left:* The four identical subunits are shown in red, yellow, green, and orange. The two substrates (dihydrofolate and NADPH) are shown in the active site (central pore). *Right:* Residues in direct contact at the dimerization and tetramerization interfaces are colored in orange and purple, respectively, along with their known dissociation constants (see Methods). Coordinates were taken from PDB: 2RK1^26^ and visualized with ChimeraX^27^. **C)** Study system in which DfrB1 complements the inhibition of FolA by trimethoprim (TMP) in *E. coli*. **D)** Experimental simulation of gene duplication using two plasmids. One plasmid contains an ampicillin resistance marker (Amp^R^) and a complete library of *dfrB1* mutations (*dfrB1*-DMS) from codon 10 onwards. The second plasmid contains a kanamycin resistance marker (Kan^R^) along with either an empty insertion site or a WT copy of *dfrB1* bearing synonymous mutations in codons 4-9 (*dfrB1[Syn4-9]*). The two copies are controlled by the same arabinose-inducible promoter.

As shown in Fig. 1, a key factor affecting the maintenance of paralogs is the fraction of LOF mutations that can interfere with the other copy. We set out to estimate this fraction using a small homomeric protein for which we can estimate the functional impact of mutations by measuring antibiotic resistance^24^. The *E. coli* plasmid-encoded DfrB1 enzyme uses NADPH to catalyze the reduction of dihydrofolate (DHF), the same reaction as the *E. coli* dihydrofolate reductase (FolA), although the two proteins are not homologous. DfrB1 forms a homotetramer whose active site is located in the central pore (Fig. 2B). While FolA is inhibited by the antibiotic trimethoprim (TMP), DfrB1 is insensitive to this molecule^25^. We monitor the impact of mutations on DfrB1 through its complementation of the growth defect caused by TMP (Fig. 2C).

We experimentally simulated a gene duplication by expressing two copies of the gene on separate plasmids while preserving their promoter and thus, their transcriptional regulation. To selectively amplify the mutated paralog for subsequent high-throughput library sequencing (see Methods, Fig. S2A), one copy of DfrB1 bears synonymous mutations in codons 4-9 (Syn4-9). We first confirmed that these mutations do not affect growth in the presence of TMP (Fig. S2B) and that protein copies from the different plasmids form heteromers (Fig. S2C). Because the two plasmids have different antibiotic markers, cells could harbor different copy numbers of each plasmid if the different number of copies of antibiotic markers were needed to achieve the same growth rate. We thus validated that expressing WT DfrB1 from the two plasmids results in a comparable growth at different induction levels (Fig. S3). Our result suggests that any variation in plasmid copy number averages out at the population level. Then, we measured the effect of all amino acid substitutions of the enzyme downstream of codon 9 while expressing a singleton mutated gene (experiment 1) or when duplicated alongside a WT copy (experiment 2) (Fig. 2D, Fig. S4A). Importantly, DfrB1 copies in both plasmids are regulated by an arabinose-inducible promoter in media with 0.0015% arabinose, the concentration required to recover 50% of the growth of cells harboring two DfrB1 copies in media without TMP (Fig. S4B). At this concentration, DfrB1 is limiting for growth, allowing the detection of both adaptive and deleterious mutations.

### Mutations causing negative paralog interference have mild destabilizing effects and cluster in structural hotspots

Globally, combining the mutant library with a WT copy led to a decrease in the magnitude of fitness effects (Fig 3A, Tables S1-S2). Most mutations that were deleterious when expressed as a singleton became milder when co-expressed with a second WT copy. We note that on the other end of the spectrum, some mutations were beneficial only as singletons. In line with a previous study^24^, we found that some of these mutations lead to an increase in DfrB1 abundance (Fig. S5, Table S3).

**Figure 3:**
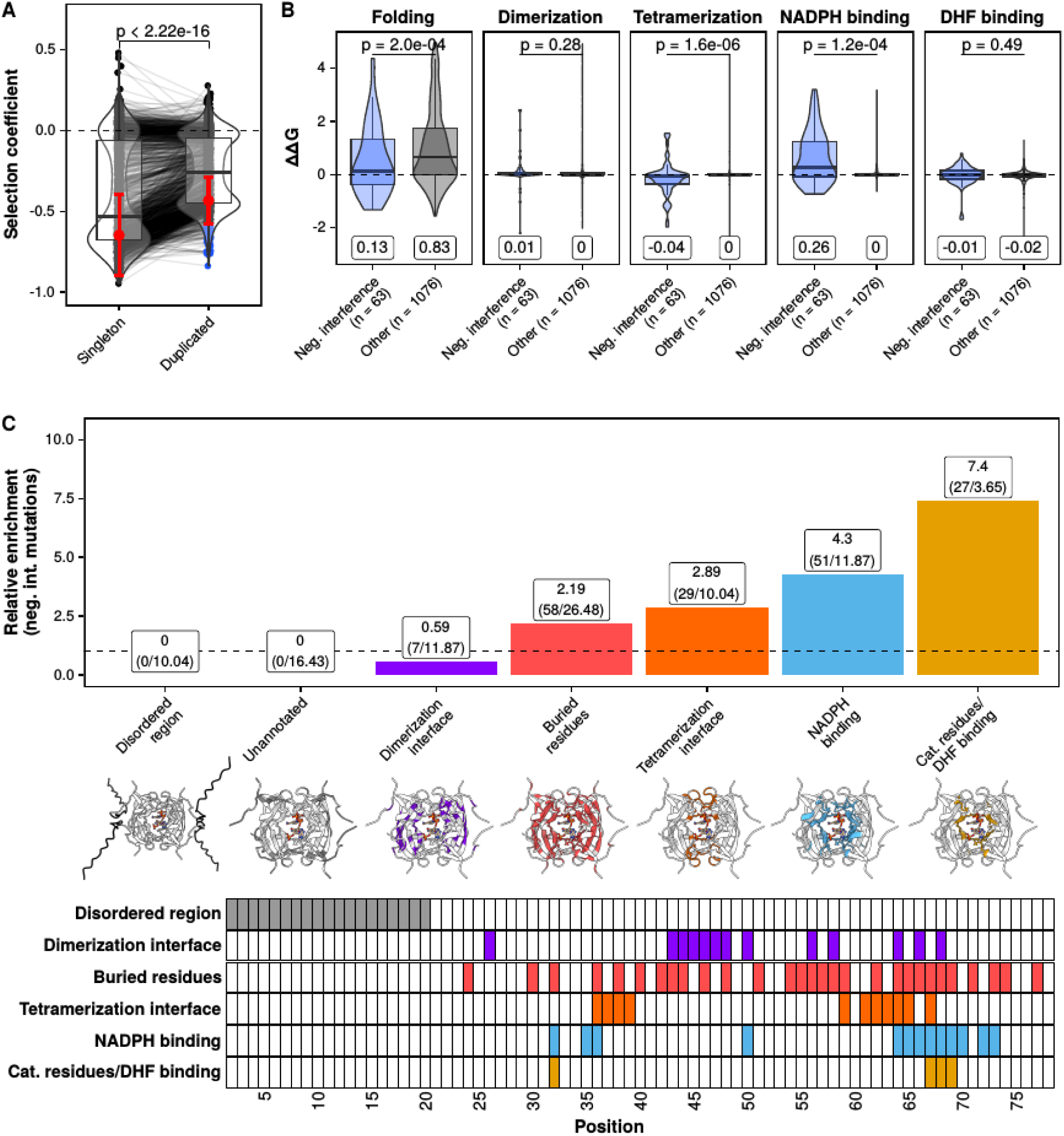
General properties of negative interference substitutions in an oligomeric complex. **A)** Distributions of measured selection coefficients for the mutant library in the singleton (n = 1443) and duplicated (n = 1448) backgrounds. Red dots and error bars represent the mean +/- 2.5 times the standard deviation of the selection coefficient of stop codons. Negative interference substitutions (blue) are conservatively defined as those that are more deleterious than the mean minus 2.5 times the standard deviation of stop codons in the duplicated background. The p-value was calculated using a paired Mann-Whitney U test. **B)** FoldX-predicted effects of negative interference substitutions on DfrB1 folding, dimerization, tetramerization, and NADPH and DHF binding compared to the effects of other substitutions. Labels under each distribution indicate their median ΔΔG values. p-values were calculated using Mann-Whitney U tests. **C)** Relative enrichment of negative interference substitutions across protein regions, calculated as a ratio of observed over expected counts. The dashed line indicates a relative enrichment of 1, in which the observed counts of negative interference substitutions are identical to the expected counts. Cartoons indicate the different protein regions tested for the enrichment of negative interference substitutions. Positions along the DfrB1 sequence belonging to each region are shown at the bottom.

Mutations causing negative paralog interference have a more deleterious fitness effect than stop codons. We initially defined high-confidence negative paralog interference candidates as mutations with fitness effects below the mean minus 2.5 standard deviations (probability of false positives = 0.006) of stop codons that are in the same competing pool with two gene copies (see Methods). In the experiment expressing the singleton mutant library, 986 out of 1310 possible amino acid substitutions (75.3%) were deleterious (worse than the 2.5 percentile of WT synonymous codons) and 766 (58.5%) were within 2.5 standard deviations of the mean of stop codons (Fig. 3A). Only two substitutions were below the lower limit of this interval, which could happen for instance if some variants were toxic or due to experimental noise. When the mutant library was expressed alongside the WT copy, 967 (73.8%) amino acid substitutions were deleterious, 554 (53.7%) behaved like stop codons, and 63 had more deleterious effects than stop codons (representing 6.4% of deleterious substitutions) (Fig. 3A, Table S4). As a result, these 63 substitutions are strongly selected against despite having a redundant WT copy of the gene present, which results from negative interference. When comparing the effects of substitutions in the singleton and duplicated backgrounds, we observed that interfering ones were not rescued by the second copy, unlike stop codons and other substitutions that became less deleterious (Fig. S6). Since the starting pool composition can affect the dynamic range of deleterious effects observable for each mutant, we compared our observed selection coefficients to the most deleterious effect observable for each variant (see Methods). Most stop codons and interfering substitutions were within the dynamic range of the assay (Fig. S7), confirming that the observed effects are not a byproduct of differences in pool composition.

We set out to identify the general properties of interfering substitutions. We computationally estimated the effects of all substitutions on the folding, dimerization, and tetramerization energies of DfrB1, as well as on the binding affinities of the DfrB1 tetramer for its substrates DHF and NADPH (see Methods, Table S2)^28,29^. Since the catalytic core of the protein is formed by the contribution of the four chains, a single destabilizing substitution in one chain of the complex could be sufficient to reduce or prevent any activity. Compared to the rest of substitutions, those causing negative interference are predicted to have milder destabilizing effects on protein folding, slightly more stabilizing effects on tetramerization affinity, and more disruptive effects on NADPH binding (Fig. 3B). We did not observe significant differences in terms of dimerization affinity and DHF binding (Fig. 3B). NADPH is the first substrate to bind in the reaction producing THF, which is consistent with its disruption being more important than that of DHF binding^30^. These results hold when comparing the interfering substitutions against others occurring at the tetramerization interface and active site (Fig. S9).

Our observations of interfering substitutions allow estimating the expected increase in residence time of paralogs with similar properties to DfrB1. Our interfering substitutions behave like LOF in our model and both stop codons and substitutions disrupting the dimerization interface by at least 2 kcal/mol correspond to LOA. Thus, out of 4116 total codons sampled for DfrB1, 180 (4.37%) encode LOF substitutions and 499 (12.12%) correspond to LOA substitutions. While we do not have an estimate of mutational effects for the arabinose promoter, we use data from mutagenesis of more than 2000 *E. coli* promoters^31^ to derive a rough estimate of excess retention times. From their data, we considered as LOE mutations those that decreased the activity of a given promoter by at least 50% (see Methods). The fraction of such mutations ranges from 0% to 68.2% across different promoters, with a median of 3.4%. Allowing the rate of LOE mutations to vary in that interval and using our estimates of LOA and LOF mutations for DfrB1, the residence time for redundant paralogs increases by 5.3%-35.8% depending on the rate of LOE mutations (Fig. S8A). In addition, we set up a Moran model to simulate the residence time of the paralogs (see Methods). Using the parameters from the previous example, we observed an increase of 8.5% in the average number of birth/death events required to reduce the number of individuals harboring redundant duplicates by 80% in a sample of 1000 populations with 1000 individuals each (Fig. S8B). Restricting the data to only 621 variants accessible by a single point mutation results in 11 (1.77%) LOF mutations and 47 (7.57%) LOA mutations. Using these parameters and the median rate of LOE mutations at 3.4%, the estimated increase in residence time due to interference corresponds to 16.1% in the analytical model (Fig. S8C) and 1.8% in the Moran model (Fig. S8D). Thus, while these estimates may vary for different proteins, our results provide an intuition regarding the order of magnitude of the effect of paralog interference in the preservation of redundancy.

Considering that negative interference substitutions have different effects on the two interaction interfaces (Fig. 3B), we asked if they were clustered in the protein structure. We calculated the relative enrichment of negative paralog interference substitutions across protein regions including the two interfaces, the active site, and buried residues (see Methods). Interfering substitutions are most strongly enriched in residues interacting with the substrates, followed by residues being located at the tetramerization interface (Fig. 3C). In turn, they are underrepresented at the dimerization interface, confirming the differences between the two interfaces. Since there is considerable overlap among protein regions, we repeated this analysis while deconvoluting the signal from buried residues, the two interfaces, and the NADPH binding site. The highest enrichment in interfering substitutions was still observed in the NADPH binding site, although the signal is stronger when it overlaps with the tetramerization interface than when it overlaps with the dimerization interface (Fig. S10). Impòrtantly, this analysis also shows that the presence of interfering mutations in buried sites results from their overlap with the tetramerization interface and the catalytic sites. The DfrB1 active site and its tetramerization interface are thus hotspots of negatively interfering substitutions.

### The magnitude of deleterious effects depends on the interference mechanism

Since interfering substitutions cluster at the catalytic site and at the tetramerization interface, but not at the dimerization interface, we set out to dissect the contributions of LOF and LOA to interference. In our initial model (Fig. 1A), we assumed that LOA substitutions do not cause interference because we considered a simple dimer. However, DfrB1 assembles in two steps as a dimer of dimers^30^. Thus, we hypothesized that substitutions disrupting the second assembly step could also sequester a fraction of WT copies in inactive heterodimers, unlike substitutions disrupting the initial, dimerization step (Fig. 4A). Importantly, such LOA alleles at the tetramerization interface would be expected to sequester a smaller fraction of WT protein copies than LOF alleles, so we hypothesized that their resulting deleterious effects would be smaller than those of LOF alleles (Fig. 4B).

**Figure 4.**
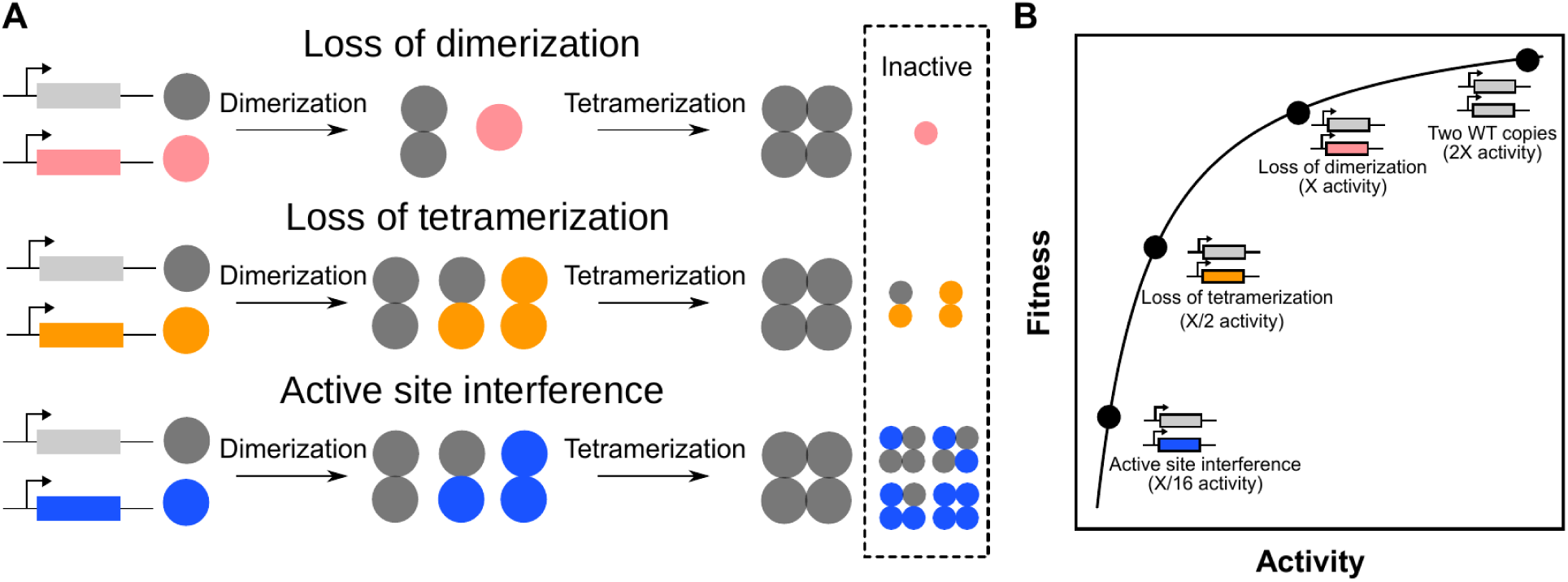
Model of paralog interference for tetramers. **A)** The loss of dimerization eliminates physical interactions and potential for interference. Mutant protein copies with disrupted tetramerization interfaces do not tetramerize and instead sequester WT copies in heterodimers. Substitutions in catalytic sites do not disrupt tetramerization, sequestering a large fraction of WT copies in inactive tetramers. **B)** Relative positions of each genotype along a fitness curve based on the concentration of active tetramers.

We selected different deleterious substitutions with predicted destabilizing effects at either of the two interfaces for further validation. In our experimental data, the only selected substitution with a clear stronger deleterious effect than stop codons was Q67C (Fig. 5A, ANOVA post-hoc HSD test p = 2.1e-5, Table S5), which has been shown to greatly reduce catalytic activity^32^. We set up pairwise competition assays (Fig. S11) for cells harboring one or two copies of DfrB1. When competed against a single WT copy, only the tetramerization interface substitution S59Y had a more deleterious effect than stop codons (Table S6, ANOVA post-hoc HSD test p = 9.9e-5), while all the others caused either intermediate deleterious effects or were not significantly different from stop codons (Fig. 5B). Nevertheless, competitions between cells expressing each variant alongside a WT copy versus cells expressing two WT copies revealed multiple interfering effects (Fig. 5C, Table S7). Substitutions disrupting the dimerization interface remained at intermediate values or equally deleterious as stop codons, showing no interfering effects.

**Figure 5.**
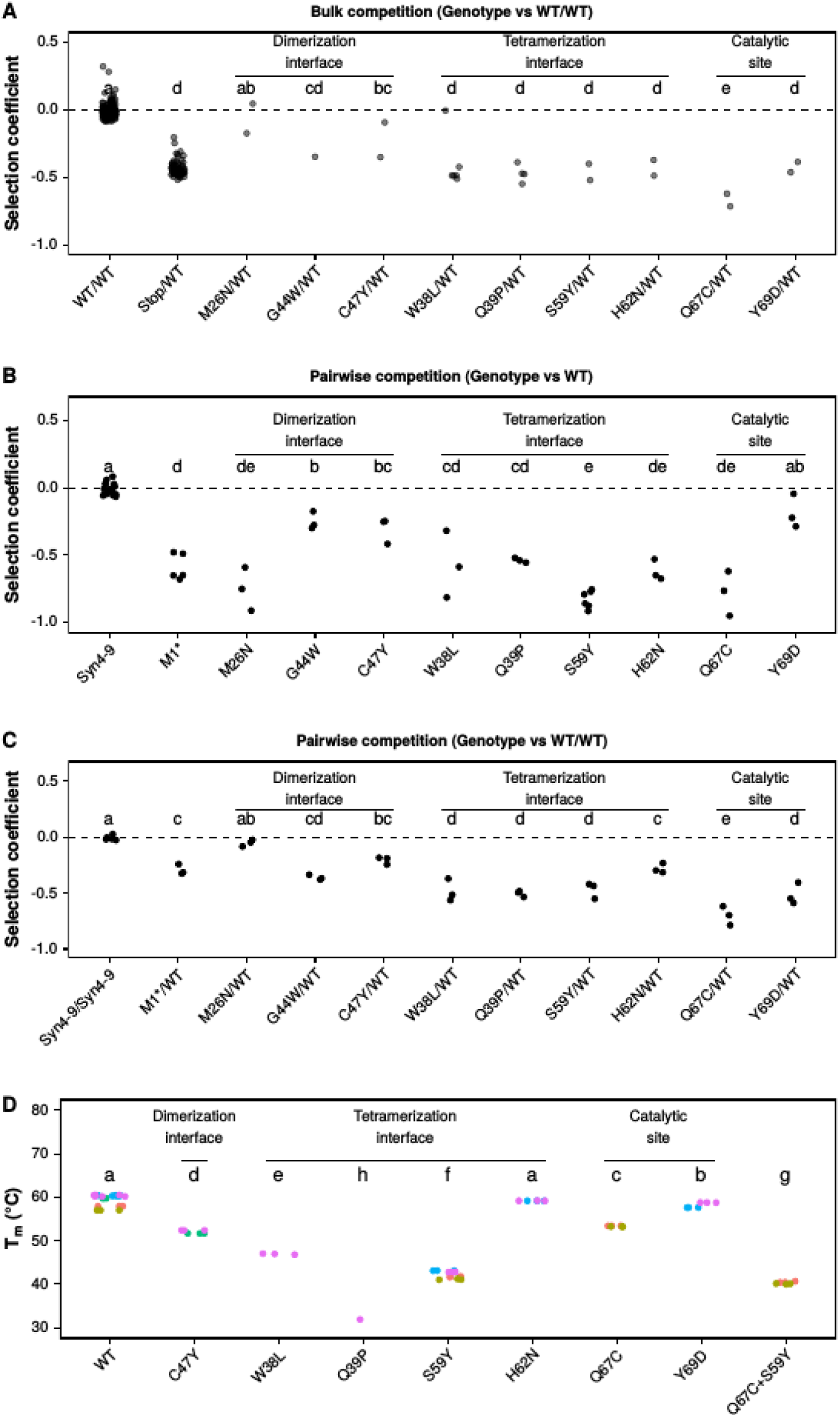
The deleterious effects of interfering substitutions vary depending on the interfering mechanism. A-C) Selection coefficients for selected mutants in different competition settings: **A)** expressed alongside a WT copy in the DMS bulk competition experiment using two WT copies (WT/WT) as a reference, **B)** as a single copy in a direct pairwise competition assay using a single WT copy as a reference, and **C)** expressed alongside a WT copy in a direct pairwise competition assay using WT/WT as a reference. Stop codons from position 10 onwards (Stop/WT) in the DMS data and at position 1 (M1*/WT) in pairwise competition assays are shown for comparison. In panels B-C, Syn4-9 refers to WT DfrB1 with synonymous mutations in codons 4-9. **D)** Temperature at which 50% of tetramers harboring different substitutions dissociate (T_m_). T_m_ was measured in triplicates for separate purifications performed on different days. Colors indicate separate purifications performed on different days. Labels at the top of panels A-D indicate significantly different groups as determined by an analysis of variance (ANOVA) and Tukey’s post hoc honestly significant difference test (p < 0.05).

Meanwhile, substitutions disrupting the tetramerization interface and the catalytic site had more deleterious effects than stop codons, capturing the effect of negative interference. These results suggest that our estimate derived from the bulk competition experiment that 6.4% of deleterious substitutions cause negative interference is likely a lower bound, dominated by LOF effects. They also support our hypothesis that the two-step assembly of the complex makes one interface more prone to negative interference than the other.

We validated the destabilizing effects of several substitutions using nano-differential scanning fluorimetry. We observed stronger destabilization from substitutions occurring at the dimer and tetramer interfaces than in the active site (Fig. 5D, Fig. S12, Table S8). These effects are in agreement with our model of the origin of their deleterious effects and their interference mechanisms. For example, substitution Y69D does not destabilize the tetramer (Fig. 5D) but still has a slightly deleterious effect in the single copy background (Fig. 5B), suggesting the mechanism is moderate LOF. This effect is amplified in the duplicated background due to interference with WT copies (Fig. 5C) and becomes indistinguishable from strong substitutions disrupting tetramerization. Similarly, despite being as deleterious as either S59Y and Q67C in singleton (Fig. S13A, Table S9), the deleterious effect of a DfrB1 copy harboring both S59Y+Q67C is more similar to S59Y than Q67C when duplicated alongside a WT copy (Fig. S13B, Table S10). In fact, S59Y+Q67C is slightly less deleterious than S59Y alone, potentially because S59Y+Q67C combined have a higher destabilizing effect on tetramerization than S59Y alone, leading to the sequestration of fewer WT copies.

Together, these results show that interfering alleles sequester functional alleles into inactive heterocomplexes either by disrupting subsequent assembly steps or catalytic activity, and that the magnitude of their deleterious effects depends on the fraction of sequestered functional subunits.

## Discussion

Gene duplication has been shown to occur at a rate roughly comparable to point mutations^1,3,8^. While previous work has suggested that homomeric proteins and genes with a high access to dominant deleterious mutations could be preferentially retained^9–12^, the underlying mechanisms remain unclear. Here, we explore how homomeric paralogs can interfere with each other, producing deleterious mutations that would impact the product of both duplicates, just as dominant mutations do within a locus. We demonstrate that selection against such interfering mutations favors the maintenance of functional redundant paralogs. Although we do not model the establishment phase prior to fixation of the duplicates, this phase would follow known principles. Neutral duplications would have a probability of fixation that is the reciprocal of the effective population size. Advantageous duplications would be more likely to fix than neutral expectation, and deleterious duplications less likely. We can assume that if negative interfering mutations were to occur before fixation of the duplicate, they would be more likely to be selected against than LOA, LOE or non-interfering LOF mutation, favoring the maintenance of functional gene copies.

Using an experimental model, we find that a lower bound of 6.4% of deleterious substitutions for a small homotetrameric protein (DfrB1) are more deleterious when expressed alongside a WT copy than entirely losing one copy, making them negatively interfering mutations. We identify two broad classes of interference mechanisms based on the detailed study of a tetrameric enzyme. First, an active site formed at the interface between subunits can be inactivated by substitutions in one subunit. Around 20% of eukaryotic protein complexes have been reported to have multi-chain ligand binding sites^33^, with some having known cases of dominant negative substitutions^34^. Second, negative interference substitutions are also prevalent at the tetramerization interface but depleted at the dimerization interface of DfrB1. Since DfrB1 assembles sequentially into dimers and then into tetramers^30^, disrupting the dimerization interface eliminates all assemblies, whereas destabilizing the tetramerization interface allows mutant alleles to sequester WT copies into non-productive heterodimers. Dimers of dimers, such as DfrB1 and human hemoglobin, are among the most common oligomeric protein complexes^35^, suggesting these negative interference mechanisms could apply to the retention of many such duplicates. Indeed, a large fraction of dominant negative mutations are found at interaction interfaces^17^. Overall, the assembly process of oligomers and the architecture of their ligand binding sites could influence the rate of interfering mutations for different proteins.

The effects of negative interference could be modulated by other mechanisms. For example, some proteins could retain some activity in their monomeric forms, as in the case of proteins showing transient and facultative interactions^36,37^. In such cases, LOA mutations would produce more monomers and be less deleterious than LOE. Nevertheless, LOF would remain the most deleterious and negative selection against them and would contribute to the preservation of redundancy. Similarly, our model assumes that all synthesized protein copies assemble into homo- or heteromers. If not all protein copies dimerize, interfering alleles would inactivate a smaller fraction of WT protein copies, reducing the magnitude of the deleterious effects of interference^38^. Conversely, interactions between protein copies have been shown to allow the accumulation of mutations that would otherwise destabilize protein folds^39,40^. Such a process could lead to the two paralogs preferentially heteromerizing or only being functional as heteromers^10,41–43^. Such pairs of paralogs could allow protein copies harboring interfering substitutions to sequester more copies of the other subunit in non-productive dimers, which would potentiate deleterious interfering effects^38^. Overall, the deleterious effects of interference and their contribution to maintaining paralog redundancy would be affected by the propensity of the paralogs to heteromerize. More experiments on other protein complexes will be necessary to examine how these different scenarios promote or not the maintenance of paralogs.

The number of genes in a genome depends on the balance between gene birth, which is dominated by gene duplication, and gene loss. The mechanisms described here by which paralogs can interfere with their sister copy alter this balance, with several implications. First, genes that are prone to negative interference, such as those encoding oligomeric proteins, should be maintained by purifying selection, even if they are redundant. As a result, their gene families could increase in size. Second, the longer retention time may help acquire sub- or neofunctionalization mutations that will eventually further enhance their retention time^7,41,44,45^. Our work therefore illustrates how mutations and biochemistry interact to shape genome content.

## Acknowledgments

We thank the Landry lab members for their discussions during the project. We thank Pascale Lemieux, François D. Rouleau, Simon Aubé, and Pavithra Venkataraman for their comments on the manuscript. We also thank Alicia Pageau and François D. Rouleau for bioinformatics support, Claudèle Lemay St. Denis and Joelle Pelletier for help with DfrB1 experiments, and Sarah Otto for comments on a previous version of our model.

## Attributions

Conceptualization: AFC, FM, YS, LNT, CRL Methodology: AFC, FM, YS, LNT, CRL Investigation: AFC, FM, IGA, YS, LNT, MMV, MCMV Experiments: FM, IGA, AKD, MMV, MCMV, AFC Visualization: AFC, FM, YS Funding acquisition: CRL Project administration: CRL Supervision: CRL Writing – original draft: AFC, CRL, FM, IGA Writing – review & editing: AFC, CRL, FM, IGA

## Data and materials availability

All raw sequencing data are available at SRA BioProject PRJNA1097537 (accession numbers SRR28586753-SRR28586760 and SRR28586807-SRR28586813). Scripts for data analyses and figure generation are available at https://github.com/Landrylab/Paralog_interference_DfrB1. All software and materials used are listed in the key resources table (Table S11).

## Materials and Methods

### Model of transitions between states

We built a model assuming transitions between four states in a haploid population: the initial state after duplication (two active copies) and the states after the acquisition of a loss-of-function (LOF), loss-of-affinity (LOA), or loss-of-expression mutation (LOE) (Fig. 1). Transitions between these states are considered to be irreversible and to only occur toward the loss of properties. The system can leave the initial state of redundancy by having one of the copies acquire LOF, LOA, or LOE mutations. Transitions between states are given by their respective mutation rates (µ_F_, µ_A_, and µ_E_) and probabilities of fixation, which vary depending on whether the model allows for paralog interference or not. The time needed to lose redundancy is inversely proportional to the probability of a transition fixing and thus follows a geometric distribution.

The expected time the system maintains redundancy is inversely proportional to the rates of these transitions, as shown in equation 1:

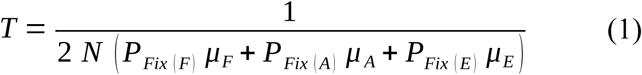

where *T* is the time needed to lose redundancy; *N* is the effective population size; *P_Fix(F)_*, *P_Fix(A)_*, and *P_Fix(E)_* are the probabilities of fixation of LOF, LOA, and LOE mutations; and *µ_F_*, *µ_A_*, and *µ_E_* are the rates of LOF, LOA, and LOE mutations.

Similarly, the variance of the time needed to lose redundancy would be given by the following expression:

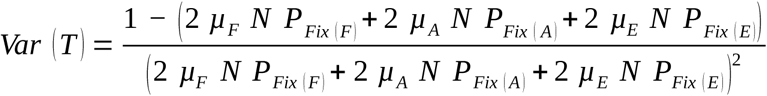

which increases as probabilities of fixation and mutation rates decrease.

### Classical model - no paralog interference

In a system without interference (Fig. 1A), mutations in one of the protein copies do not affect the function of the other copy. Thus, the probabilities of fixation in equation 1 for LOF, LOA, and LOE mutations are then identical and can be replaced by a general term (*P_Fix(Loss)_*) that reflects the loss of activity of one copy, as shown in equation 2:

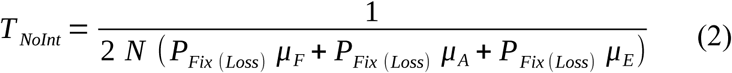

### Model with paralog interference

As opposed to the classical model, mutations in one copy can modify the function of the second copy in a system with paralog interference (Fig. 1B). Since such modifications rely on the physical interaction between the two protein copies, they are only possible for LOF mutations and not for LOA and LOE mutations. As a result, the probabilities of fixation for LOA and LOE mutations are identical (*P_Fix(Loss)_*), but different from that of LOF mutations (*P_Fix(F)_*). equation 1 can then be simplified to obtain the general form of the model with interference, as shown in equation 3:

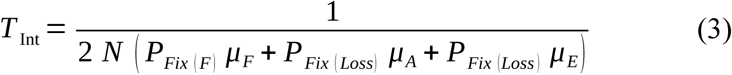

### Comparison of time needed to lose redundancy assuming neutrality

In many cases the presence of a duplicate gene makes the loss of the second copy neutral^5,46^. In such a scenario, one can assume that LOF, LOA, and LOE mutations in one copy are neutral, and thus *P_Fix(Loss)_ = 1/N*. As a result, this assumption allows simplifying the equations of the classical and interference models to obtain equations 4 and 5, respectively:

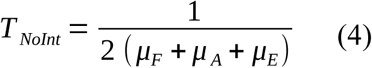

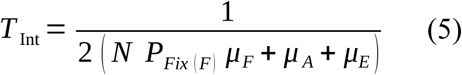

The times needed to lose redundancy for the two models can be compared by using the ratio shown in equation 6:

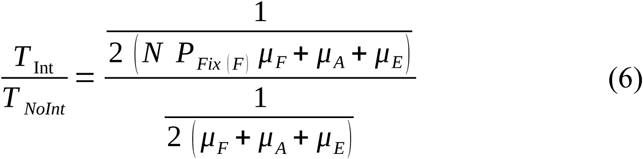

After applying the division:

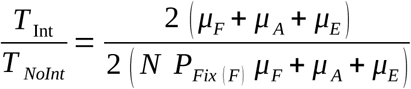

After cancelling terms and dividing by *µ_F_*:

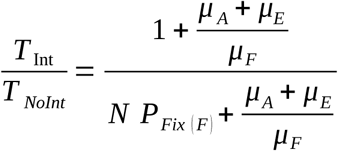

This expression allows comparing the times needed to lose redundancy in both models. To identify the conditions in which paralog interference causes the system to maintain redundancy for longer, we can use the following inequality:

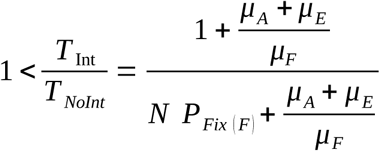

After multiplying both sides by the denominator of the right-hand side:

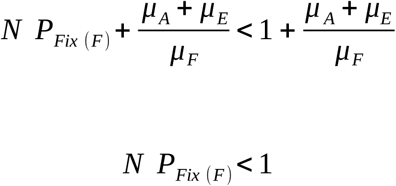

This expression shows that the time needed to lose redundancy is higher in the model with interference if *NP_Fix(F)_ < 1*. Since *P_Fix(Loss)_ = 1/N*, the requirement needed to preserve redundancy for longer in the system with interference is equivalent to *P_Fix(F)_ < P_Fix(Loss)_*.

### Comparison of times needed to lose redundancy without the assumption of neutrality

Alternatively, the loss of a gene copy after the duplication may not be neutral. Our model can also be used to compare the times needed to lose redundancy even when this assumption is relaxed, as shown in equation 7:

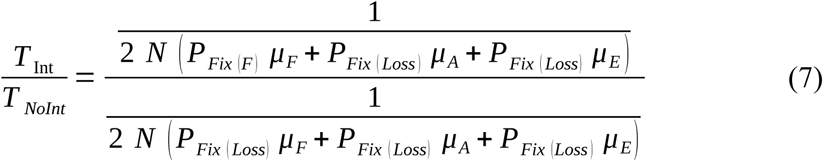

After applying the division:

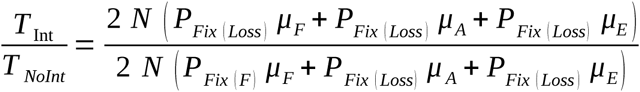

After cancelling terms:

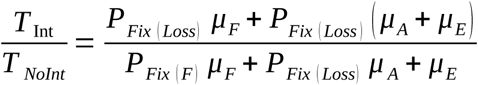

Dividing by *µ_F_*to show the expression in terms of *(µ_A_ + µ_E_) / µ_F_*:

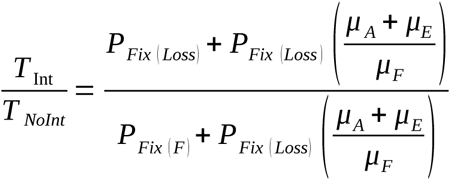

This expression allows comparing the times needed to lose redundancy in both models. To identify the conditions in which paralog interference preserves redundancy for longer, we can use the following inequality:

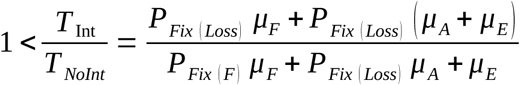

After multiplying both sides by the denominator of the right-hand side:

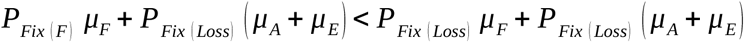

After cancelling terms:

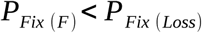

This expression shows that the time needed to lose redundancy is greater in the model with interference if *P_Fix(F)_ < P_Fix(Loss)_*. The probability of fixation (*P_Fix_*) of a mutation depends on its selection coefficient (*s*). Assuming small fitness effects, the probability of fixation is on the order of *s* for haploid cells and 2*s* for diploid cells^47^. Thus, the condition for redundancy to be preserved for longer in the model with interference becomes:

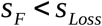

Or, in other words, that the interfering LOF mutations are more deleterious than the loss of one copy.

### Strains and media

The *Escherichia coli* strains MC1061 and BL21(DE3) were used for cloning and mutagenesis while only BL21 (DE3) was used to conduct all the experiments in this study. All transformations with plasmids were done with in-house protocols for chemocompetent cells. Transformed cells were grown on selective 2xYT + glucose medium (1.0% yeast extract, 1.6% tryptone, 0.2% glucose, 0.5% NaCl, and 2% agar) with either ampicillin (AMP; 100 μg/mL), kanamycin (KAN; 50 μg/mL), or both. For all experiments conducted in liquid medium, bacteria were grown in Luria-Bertani (LB) medium (0.5% yeast extract, 1.0% tryptone, and 1.0% NaCl) with the corresponding antibiotics (AMP; KAN; or AMP + KAN) and with or without L(+)-arabinose (Ara) and trimethoprim [TMP −10 μg/mL in dimethyl sulfoxide (DMSO)]. Protein induction for purification was achieved in Terrific Broth (TB) medium (2.4% yeast extract, 1.2% tryptone, 0.94% K_2_HPO_4_, 0.22% KH_2_PO_4_, and 0.4% glycerol).

### Plasmid construction

All of the following PCR amplifications were done using KAPA HiFi HotStart polymerase unless specified otherwise. Oligonucleotide and gBlock sequences and descriptions used in this study can be found in Table S12. All plasmids were recovered from bacteria using the Presto^TM^ Mini Plasmid Kit. The integrity of assembled and mutagenized plasmids was confirmed by Sanger sequencing (Plateforme de séquençage et de génotypage des génomes, Centre de recherche du centre hospitalier de Québec–Université Laval (CRCHUL), Canada).

For the bulk competition experiments, *dfrB1* gene was expressed in bacteria from a pBAD vector that allows arabinose-controlled induction^48^. In addition to the same pBAD-*dfrB1* (that we will hereafter call pBAD-*dfrB1*-AMP to distinguish from the one that has a KAN resistance module) that was used by^24^, minor modifications of the vector were also made for this study. First, the ampicillin resistance cassette was replaced by a kanamycin resistance cassette to generate pBAD-*dfrB1*-KAN. For this, the pBAD-*dfrB1*-AMP plasmid was amplified by PCR (PCR program: 5 min at 95°C; 35 cycles: 20 s at 98°C, 15 s at 65°C, and 2 min at 72°C; and a final extension of 3 min at 72°C), removing the ampicillin marker. The KANMX resistance cassette was amplified from pMoBY-ACT1 (PCR program: 5 min at 95°C; 5 cycles: 20 s at 98°C, 15 s at 61°C, and 1 min at 72°C; 30 cycles: 20 s at 98°C and 1 min 15 s at 72°C; and a final extension of 3 min at 72°C). Both PCR products were incubated with 20 U of DpnI for 1 hour at 37°C to remove template DNA, and subsequently purified on magnetic beads following the manufacturer’s instructions. The plasmid (pBAD-*dfrB1*) and insert (KANMX), with their overlapping regions at each extremity, were then assembled by Gibson DNA cloning^49^, producing pBAD-*dfrB1*-KAN plasmid.

The pBAD-*dfrB1*-KAN plasmid was then used to generate two additional plasmids, one removing *dfrB1* (pBAD-EMPTY-KAN) as a control for the effect of duplication, and another introducing synonymous mutations from position 4 to 9 in the coding sequence of *dfrB1* (pBAD-*dfrB1[Syn4-9]*-KAN) to selectively amplify each *dfrB1* copy in subsequent high throughput sequencing library preparation. Plasmid pBAD-EMPTY-KAN was constructed by amplification of pBAD-*dfrB1*-KAN with oligonucleotides that excluded *dfrB1* coding sequence (PCR program: 5 min at 95°C; 35 cycles: 20 s at 98°C, 15 s at 68°C, and 2 min 30 s at 72°C; and a final extension of 8 min at 72°C). The PCR product was incubated with 20 U of DpnI for 1 hour at 37°C to remove template DNA, and directly transformed into BL21 competent cells for re-circularization, producing pBAD-EMPTY-KAN plasmid. Plasmid pBAD-*dfrB1[Syn4-9]*-KAN was constructed by amplifying pBAD-*dfrB1*-KAN with oligonucleotides that allow replacement of codons 4-9 in *dfrB1* coding sequence (PCR program: 5 min at 95°C; 35 cycles: 20 s at 98°C, 15 s at 61°C, and 2 min 40 s at 72°C; and a final extension of 8 min at 72°C). The PCR product was incubated with 20 U of DpnI for 1 hour at 37°C to remove template DNA, and subsequently purified on magnetic beads. Then, 100 ng of the purified PCR product was incubated with T4 polynucleotide kinase for 30 minutes at 37°C in T4 ligase buffer to perform the 5’ phosphorylation, then cooled down on ice for 3 minutes and directly incubated with 10 U of T4 ligase for 1 hour at 37°C. Half of the reaction was then transformed into BL21 cells producing pBAD-*dfrB1[Syn4-9]*-KAN. Often used as a control in many experiments, pBAD-*dfrB1[Syn4-9]*-AMP plasmid was also constructed the same way starting from pBAD-*dfrB1*-AMP.

To perform co-immunoprecipitations of the heteromers, we generated vectors to express 6xHIS and FLAG N-terminal tagged DfrB1 proteins. We amplified the backbone of the desired plasmids (either pBAD-*dfrB1*-AMP or pBAD-*dfrB1[Syn-4-9]*-KAN) with the appropriate primers depending on whether the plasmid was bearing the native or the Syn4-9 sequence *dfrB1*, respectively. The resulting PCR product was digested with 20 U of DpnI and purified using magnetic beads as described previously. Then, 50 ng of the purified products and 10 ng of the desired gBlock containing 6xHIS- or FLAG-tag sequence were assembled by Gibson DNA assembly^49^ and transformed into BL21 competent cells to retrieve the following plasmids: pBAD-6xHIS-*dfrB1*-AMP, pBAD-FLAG-*dfrB1*-AMP, pBAD-6xHIS-*dfrB1[Syn-4-9]*-KAN and pBAD-FLAG-*dfrB1[Syn-4-9]*-KAN.

In order to generate the plasmids needed to measure protein abundance of different mutants of DfrB1, we performed site-directed mutagenesis based on the QuickChange Site-Directed Mutagenesis System (Stratagene, USA). Briefly, we amplified the pBAD-*dfrB1-sfGFP* plasmid using pairs of primers containing the desired mutation at the center (PCR program: 2 min at 95°C; 22 cycles: 20 s at 98°C, 15 s at 68°C, and 4 min at 72°C; and a final extension of 5 min at 72°C). The PCR products were then incubated for 1 hour at 37°C with 20 U of DpnI enzyme to remove parental DNA, and mutated plasmids were retrieved directly by transformation in MC1061 bacteria. We used this method to generate the following mutants: pBAD-*dfrB1*(N15D)-*sfGFP*, pBAD-*dfrB1*(F18W)-*sfGFP*, and pBAD-*dfrB1*(S20M)*sfGFP*.

For validation of the bulk competition experiment fitness results, some new plasmids were needed. To produce pBAD-*dfrB1[Syn4-9]*(Q67C)-KAN, the mutant pBAD-*dfrB1*(Q67C)-AMP was isolated from the DMS library used in^24^ and Syn4-9 mutations and KANMX resistance cassette were introduced as described above. Mutants with the M26N, G44W, C47Y, W38L, Q39P, S59Y, H62N and Y69D substitutions or with a stop codon (TGA) at position 1 (M1*) to prevent translation were also generated by site-directed mutagenesis on pBAD-*dfrB1[Syn4-9]*-KAN as was done to introduce Syn4-9 mutations, except for C47Y which was generated as specified for pBAD-*dfrB1[Syn4-9]*-KAN.

To evaluate the effect of mutations on DfrB1 stability, proteins were expressed from pQE32 (Qiagen, USA) derived plasmids. Starting from pQE32-*dfrB1*, mutations C47Y, W38L, Q39P, S59Y, H62N, Q67C and Y69D were introduced by site-directed mutagenesis as explained above for mutations introduced in pBAD-*dfrB1-sfGFP* except for pQE32-*dfrB1*(C47Y) that was generated as specified for pBAD-*dfrB1[Syn4-9]*-KAN. A second round of mutagenesis was needed to introduce Q67C in pQE32-*dfrB1*(S59Y). At the end, all corresponding pQE32-*dfrB1* plasmids were generated.

To measure the effect of dominant negative mutations on the fitness of cells expressing DfrB1, we used several constructions made in our arabinose-inducible system (pBAD-*dfrB1*-AMP, pBAD-*dfrB1*(Q67C)-AMP, pBAD-*dfrB1*(S59Y)-AMP and pBAD-*dfrB1*-KAN). A last additional construction was needed to express DfrB1 Q67C and S59Y double mutant. pBAD-*dfrB1*(Q67C+S59Y)-AMP was generated by site-directed mutagenesis as explained above, starting from pBAD-*dfrB1*(S59Y)-AMP and introducing Q67C substitution.

### Co-immunoprecipitations of heteromers

The plasmids allowing expression of the 6xHIS- or FLAG-tagged proteins were transformed in various combinations into BL21 competent cells. After protein induction, total proteins were extracted using NEBExpress *E.coli* Lysis Reagent as specified by the manufacturer. The soluble protein fraction was then equally divided for co-immunoprecipitation with either Dynabeads™ His-Tag Isolation and Pulldown or ANTI-FLAG^®^ M2 Affinity Gel following the procedure established by the manufacturers. The purified proteins were migrated on an SDS-PAGE gel and detected by western blotting using THE^TM^ Anti Flag-Tag as a first antibody and an IRDye 800CW Goat anti-Mouse IgG as a secondary antibody^50^.

### Bulk competition experiments and library sequencing

We performed bulk competition experiments as previously described^24^ with slight modifications. First, we transformed BL21 with either pBAD-EMPTY-KAN or pBAD-*dfrB1[Syn4-9]*-KAN. Then, each of the strains was transformed with 75 ng of each position mutant pool from a single-site mutation library of *dfrB1* (pBAD-*dfrB1*-AMP DMS library) generated previously by a PCR-based saturation mutagenesis method^24^. Positions 1 to 10 were omitted because they would not be captured by our sequencing approach. These positions are in the N-terminal disordered region of the protein and have been shown to be largely insensitive to mutations. Transformation efficiency was estimated by counting the colonies on each plate. A yield of >640 colonies for each position was needed to ensure a minimum of a 10x coverage for each codon mutant. When the desired transformation efficiency was achieved, all the colonies on the plates were resuspended in 5 mL of 2xYT medium, OD_600_ was measured for each pool, and 15% glycerol was added to the medium to store each pool individually at −80°C for further use. In parallel, pools were equally mixed at a total OD_600_=25 (OD_600_ of individual pools were adjusted to ∼0.36) to generate a starting mutant master pool for the bulk competition assay (Fig. S4A).

Bulk competition assays were performed by first inoculating 5 mL of LB AMP+KAN medium with the mutant master pool (in either the background without (pBAD-EMPTY-KAN) or with (pBAD-*dfrB1[Syn4-9]*-KAN) an additional copy of DfrB1) to an OD_600_ = 0.01 and grown overnight at 37°C under agitation (250 rpm). Cultures were then diluted 1:100 into 5 mL of LB AMP+KAN containing 0.0015% arabinose (LB AMP+KAN+Ara) to induce the expression of *dfrB1* to a level at which the culture reaches 50% of growth recovery in the presence of TMP (see Bacterial growth curves section below, Fig. S2B). After 18 hours of incubation at 37°C under agitation (250 rpm), 3 mL of the culture was used to extract plasmid DNA of the initial timepoint (t0). In parallel, three biological replicates were generated by diluting the culture to an OD_600_ = 0.025 into 5 mL of LB AMP+KAN+Ara+TMP. The cells were incubated at 37°C under agitation (250 rpm) until an OD_600_ of 0.8 was reached (roughly five generations). The cultures were diluted 1:40 again into 5 mL of fresh LB AMP+KAN+Ara+TMP and incubated at 37°C under agitation (250 rpm) until an OD_600_ of 0.8 was reached again. At this step, 3 mL of the culture were used to extract plasmid DNA of the final timepoint (t10). At both timepoints (t0 and t10), glycerol stocks were generated by adding glycerol to a final concentration of 15% and kept at −80°C.

To ensure that only *dfrB1* sequence from the DMS library and not from the duplicated paralog would be amplified, PCR specificity was first tested using primers targeting WT or synonymous DfrB1 sequence at codons 4-9 in PCR reactions with either pBAD-*dfrB1[Syn4-9]*-AMP or pBAD-*dfrB1*-KAN as template (Fig. S2A) (PCR program: 5 min at 95°C; 35 cycles: 20 s at 98°C, 15 s at 59°C, and 20 s at 72°C; and a final extension of 2 min at 72°C). Then, plasmid DNA from t0 and t10 was used for high-throughput DNA sequencing library preparation that involved two PCR steps. A first PCR was performed to amplify *dfrB1* from the DMS library and to add the primer binding sites for the Illumina i5 and i7 Nextera indexes (Table S13). This PCR was done using as template 1 ng of the plasmid DNA for t0 and t10 obtained from the bulk competition assay using primers with a variable number of Ns in the sequence (from 0 to 3) to generate diversity in the amplicon pool (PCR program: 3 min at 98°C; 20 cycles: 30 s at 98°C, 15 s at 60°C, and 30 s at 72°C; and a final extension of 1 min at 72°C). The PCR product was verified on a 0.8% agarose gel. Then a second PCR was performed using primers to add the i5 and i7 Nextera Illumina indexes for automatic demultiplexing at the end of the sequencing process. This PCR was done using 1 μL of a 1:1000 dilution of the first PCR as a template (PCR program: 3 min at 98°C; 18 cycles: 30 s at 98°C, 15 s at 61°C, and 35 s at 72°C; and a final extension of 1 min at 72°C). Each reaction for the second PCR was performed in quadruplicate and then combined. Each sample was amplified with a different combination of primers (Table S13). Finally, the PCR product was verified on a 0.8% agarose gel, purified using magnetic beads, and sent for paired-end 250-bp sequencing on an Illumina NovaSeq 6000 (Plateforme de séquençage et de génotypage des génomes, CRCHUL, Canada).

### Sequencing data analysis

The quality of the sequencing data was analyzed using FastQC version 0.11.4^51^ and the paired reads were merged with Pandaseq version 1.1.5^52^. Cutadapt version 4.2^53^ was used to trim the reads by detecting adapters preceding and following positions 10 and 78 respectively using the following argument (*-a AGTAGCAATGAAGTCAGT…GTTTAAACGGTCTCCAGC*) to retain only the range of positions on which DMS was performed. The trimmed identical reads were aggregated using vsearch 2.15.1^54^ and aligned to the reference sequence of *dfrB1* using EMBOSS Needle version 6.6.0.0^55^ to identify the mutant codons in each read. Mutations to stop codon TAG were not considered in the data analysis, since this stop codon has been shown to have a lower termination efficiency in *E. coli*^56,57^. Stop codons at positions 15 and 78 were also excluded because they were outliers with high fitness. Variants were counted for each sample and normalized by the total number of reads in the library. The dataset was filtered to remove all the variants with <100 counts at t0 to ensure a high sensitivity and limited bias for our selection coefficient calculations. The resulting filtered read proportions were used to calculate selection coefficients based on changes in frequency for each variant between the starting and the end point of the bulk competition assay normalized to the median of WT. The same equation was used as in Cisneros et al.^24^:

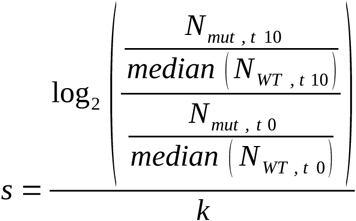

where *s* is the selection coefficient, *N* is the number of reads for the corresponding mutant at a specific time point, and *k* is the number of generations (*k* = 10). The final selection coefficients were the average of those obtained from three replicates. Selection coefficients for amino acid substitutions were calculated as the median of the selection coefficients of all corresponding synonymous variants (Table S2). The most deleterious observable selection coefficients were calculated using the same formula but setting the final read count to zero and adding a pseudocount of one.

### Identification of negative interference mutations

Negative interference amino acid substitutions (Table S4) were defined as those that had a fitness below the mean of nonsense mutations minus 2.5 standard deviations in the experiment with the duplicated DfrB1 background. Selection coefficients were calculated as indicated above for all missense and nonsense mutations. Substitutions were compared to the mean of nonsense mutations and visualized in Fig. 3A-B.

The enrichment of negative interference substitutions was calculated as the ratio of their observed over expected counts. To calculate the expected numbers of negative interference substitutions for each protein region, we first calculated the overall percentage of negative interference substitutions by dividing their total count over the total number of possible substitutions for the whole protein. Then, we multiplied this overall percentage by the number of possible substitutions in each region (the number of positions times 19). The resulting number was the expected number of negative interference substitutions for each protein region.

### Estimation of the extension of residence times for DfrB1

#### Analytical model

Our model from the previous sections requires rates of LOF, LOA, and LOE mutations. The rates of LOF and LOA mutations were obtained based on the 4116 total codons sampled for DfrB1.

LOF mutations corresponded to the 180 (4.37%) mutations encoding interfering substitutions, whereas LOA mutations were the 499 (12.12%) mutations encoding stop codons or substitutions that destabilize the dimerization interface by at least 2 kcal/mol. Since we do not have direct data for the arabinose promoter, the rates of LOE mutations were obtained from a previous analysis of more than 2000 promoters^31^. Briefly, that study divided each promoter in 10 bp fragments and scrambled the sequences of each fragment one at a time. We considered the rate of LOE mutations as the proportion of scrambled fragments that result in a decrease in expression of at least 50%. Across 2034 promoters with at least 15 scrambled fragments, the proportion of LOE mutations ranges from 0% to 68.2%, with a median of 3.4%. Setting the rates of LOF, LOA, and LOE mutations respectively to 4.37%, 12.12%, and 3.4%, we obtain the ratio (µ_E_+µ_A_)/µ_F_=3.56.

Assuming that the loss of one copy is neutral, the corresponding values were then plugged in equation 6. The probability of fixation for LOF mutations was calculated using the average selection coefficient observed for interfering mutations (s = −0.64) and the following equation^47^:

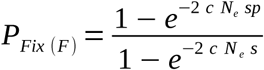

where *P_Fix(F)_*is the probability of fixation for LOF mutations, *c* is the ploidy of the organism (1 for a tandem gene duplication in a haploid genome), *N_e_*is the effective population size (set to 1e6), *s* is the selection coefficient of LOF mutations, and *p* is the number of individuals in which the duplication is introduced (set to *p* = 1/*N_e_*, with the assumption that the effective population size equals the census population size).

The resulting curve (Fig. S8A) shows the range of ratios of time needed to lose redundancy with and without interference for different rates of LOE mutations. We also repeated this analysis when considering only mutations accessible by a single point mutation from the WT sequence. This condition restricted our analysis to 621 accessible variants, comprising 11 (1.77%) LOF mutations and 47 (7.57%) LOA mutations. The corresponding curve with varying rates of LOE mutations is shown in Fig. S8C. Setting the rates of LOF, LOA, and LOE mutations respectively to 1.77%, 7.57%, and 3.4%, we obtain the ratio (µ_E_+µ_A_)/µ_F_=6.22.

#### Moran model

We set up a Moran model with selection following the description by Otto and Day^58^ to simulate the loss of redundancy between paralogs across individuals in a population. Briefly, we start with a population of N = 1000 individuals. At every step, an individual is chosen at random to become a parent for a new individual. The probabilities of selecting a parent with a given genotype are based on the frequency of that genotype in the population and the fitness of that genotype (1 for the WT, 1 + *s* for other genotypes). Selection coefficients for LOF mutations are set to *s* = −0.64 in the case with interference, while in the case without interference the selection coefficients for LOF, LOA, and LOE mutations are all *s* = 0. The new individual inherits the same genotype of its parent unless a mutation occurs. The probabilities of LOF and LOA are set to the same rates as above, with the rate of LOE mutations corresponding 3.4%, the median of the 2034 promoters with available data^31^. Then, the new individual replaces one of the remaining N-1 individuals, selected at random. We do not weigh probabilities by the selection coefficients at the death step to avoid applying natural selection twice. Using this model, we sample individuals until the frequency of individuals with redundant copies reaches 0.2N. The distributions of times required for 80% of individuals to lose redundancy with all sampled mutations (Fig. S8B) and only accessible mutations (Fig. S8D) are calculated by repeating the simulations with samples of 1000 populations with and without interference.

### Protein abundance measurements by flow cytometry

We use the measure of GFP level by cytometry to approximate DfrB1 abundance. A first preculture in 2 mL LB AMP medium was grown with bacteria containing the plasmid of interest (pBAD-*sfGFP* (negative control), pBAD-*dfrB1*-*sfGFP*, pBAD-*dfrB1*(N15D)-*sfGFP*, pBAD-*dfrB1*(F18W)-*sfGFP*, pBAD-*dfrB1*(S20M)-*sfGFP* and pBAD-*dfrB1*(E2R)-*sfGFP* (positive control)). As for the bulk competition assay, after an overnight incubation at 37°C with agitation (250 rpm), cells were diluted 1:100 in 2 mL of fresh medium with the addition or not of 0.2% arabinose. Following an 18-hour incubation at 37°C (250 rpm), cells were diluted again at OD_600_ = 0.025 in a final volume of 2 mL of fresh medium again containing or not arabinose. Cultures were then incubated as above for 2 hours. At this final time point, small aliquots of cells were taken and diluted in sterile filtered water to an OD_600_ = 0.05 in 200 μl. GFP fluorescent measurements and forward scatter (FSC) and side scatter (SSC) data were collected from a Guava easyCyte HT cytometer (Cytek^®^, USA). From the cytometry data, *E. coli* cells were selected on the basis of FSC and SSC. From the selected data points, the GFP fluorescence signal was measured after excitation with a blue laser (wavelength, 488 nm) and detection in the green channel (525/30 nm). The experiment was run in triplicate.

### Direct pairwise competition assays between mutants

For validation of the bulk competition experiment fitness results, we performed a direct competition assay using a modified quantitative PCR method for fitness measurements relative to a reference competitor^59–62^ (Fig. S11A). Two kinds of assays were performed. For the single copy assay, each competitor bore one plasmid coding for either the WT DfrB1 (ampicillin marker) or for different substitutions of DfrB1 (M1*, M26N, G44W, C47Y, W38L, Q39P, S59Y, H62, Q67C or Y69D - kanamycin marker, Syn4-9 mutations). For the duplication competition, each competitor bore this time two plasmids, conferring resistance to both ampicillin and kanamycin. In total ten genotypes (M1*/WT, M26N/WT, G44W/WT, C47Y/WT, W38L/WT, Q39P/WT, S59Y/WT, H62/WT, Q67C/WT or Y69D/WT)) were tested in pairwise competition against WT/WT duplication. In each pairwise competition experiment, the reference competitor harbored two copies of WT *dfrB1* (WT/WT), while the selected genotypes (mutant/WT) carried the synonymous Syn4-9 mutations in both copies. First, all competitors were grown overnight individually in a selective LB AMP or KAN medium with 0.0015% arabinose at 37°C under agitation (250 rpm). For the grown cultures of each competitor, OD_600_ was measured, adjusted to OD_600_ = 1 and then combined at a 1:1 ratio in 500 μL. A 100 μL aliquot was kept at 4°C for using directly 1 μL in the downstream qPCR. The combined competitors were then diluted to an initial OD_600_ = 0.025 in LB AMP+KAN medium with 0.0015% arabinose and TMP and incubated for 18 hours at 37°C under agitation (250 rpm). Final OD_600_ was measured to calculate the number of generations that had passed and 1 μL of the culture was directly used to quantify the replication of each competitor by quantitative PCR (QuantStudio 6 Pro Real-Time System (Thermo Fisher Scientific, USA)) using Fast SYBR™ Green Master Mix and primers specific to pBAD-*dfrB1* or pBAD-*dfrB1[Syn4-9]*^63^. A standard curve was performed using 5-fold dilutions of purified plasmid of pBAD-*dfrB1*-AMP and pBAD-*dfrB1[Syn4-9]*-AMP (Fig. S11B). All direct pairwise competition experiments were performed in triplicate.

Selection coefficients for each competitor relative to the other were calculated using a similar equation to the one used above in the bulk competition experiment:

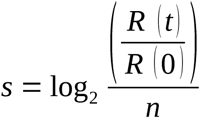

where R(0) and R(t) represent the ratio of the competitor 1 to competitor 2 in the initial (t = 0) mixture used for the competition and after 18 hours of competition (t), respectively. n represents the number of generations calculated as n=log2[OD_600_(t)/OD_600_(0)].

### Estimation of biophysical effects of substitutions

The biophysical effects of substitutions on DfrB1 folding, dimerization, and tetramerization were taken from^24^. Briefly, we used FoldX version 5.0^29^ with the MutateX workflow^64^ on the biological assembly of DfrB1 (PDB: 2RK1)^26^. Two residues from different subunits are identified as contacting if they are within a distance smaller or equal to the sum of van der Waals radii of the two closest atoms + 0.5 (Fig. 2B).

The biological assembly for the DfrB1 tetramer structure with DHF and NADPH (PDB: 2RK1)^26^ was downloaded from the PDB^65^. Since the reference structure contains an incomplete DHF molecule, the complete DHF molecule (ZINC000004228265) was downloaded from the Zinc20 database^66^ and superimposed to the coordinates in the reference PDB structure to replace the incomplete molecule. The DHF and NADPH molecules were then parameterized using Obabel version 3.1.0^67^ and the Rosetta molfile_to_params.py script^68^. The effects of substitutions on DHF and NADPH binding affinity were simulated with the Rosetta flexddG protocol^69^ using the ref2015 energy function^70^ as described previously^71^. Briefly, for each substitution 35 structures were generated and relaxed with 35 000 backrub steps following a previously described workflow^72^. From the resulting distribution of structures, the median *ΔΔG* values for mutational effects were used for all subsequent analyses.

### Protein expression, purification and stability assessment

To evaluate the effect of substitutions on DfrB1 stability, we first purify a 6xHIS-tagged version of DfrB1 and of its mutants (expression from pQE32 plasmid) as is done in^41^ with the following modifications. *E. coli* BL21 containing plasmid pRep4 was used for expression. Induction was done for 3 hours at 37°C instead of at 16°C and without zinc acetate. Cell pellets equivalent to 50 mL of post-induction cell culture were used for extraction in 2 mL of NEBExpress® E. coli Lysis Reagent. Elution buffer was finally exchanged for potassium phosphate 50 mM pH 8.0. The resulting purified DfrB1 proteins retain their N-terminal 6xHIS-tag, and were used fresh for stability assessment. All of the purification process was done in duplicate.

Protein complex stability was first measured using nano-differential scanning fluorimetry on a Prometheus NT.48 (NanoTemper, Germany). After purification, 10 μL of protein samples were loaded in high sensitivity capillaries. Measurements were taken along a 20°C to 95°C temperature gradient, increasing at a rate of 1°C/min and at an intensity of 20%. The Tm was measured by finding the temperature at which the first derivative of the 350/330 nm fluorescence ratio is maximized using the PR. ThermControl v2.3.1 (Fig. S12A). Each protein variant was tested in triplicate for both purification samples.

DfrB1 being a tetramer, in order to assess the impact of substitutions on complex assembly, 6 mg of purified protein samples were also loaded on a NativePAGE™ Bis-Tris Mini Protein Gel, 4 to 16% along with 10 μL of NativeMark^TM^ Unstained Protein Standard. Gel was run for 65 min at 300V on ice according to manufacturer’s protocol. Gel was unstained in water for 24 hours (Fig. S12B).

### Bacterial growth curves

Individual growth curves were used in four different contexts: 1) To make sure there is no fitness difference when expressing *dfrB1* in comparison to *dfrB1[Syn4-9]*, 2) To test if WT DfrB1 expressed from either of the two plasmids resulted in the same growth recovery, 3) To evaluate the concentration of arabinose needed to express *dfrB1* in duplication such as growth recovery is around 50% in presence of TMP, and 4) to validate the effect on fitness of dominant negative mutations.

In all these cases, to measure fitness for individual WT or mutant DfrB1 in duplication with a WT copy or as a singleton, growth of bacteria was followed with serial OD_600_ measurements in a plate reader. Bacteria were first transformed with 1) pBAD-*dfrB1*-AMP or pBAD-*dfrB1[Syn4-9]*-AMP, 2) pBAD-*dfrB1*-AMP or pBAD-*dfrB1*-KAN, 3) pBAD-*dfrB1*-AMP and pBAD-*dfrB1*-KAN, and 4) pBAD-*dfrB1*-AMP, pBAD-*dfrB1*(Q67C)-AMP, pBAD-*dfrB1*(S59Y)-AMP or pBAD-*dfrB1*(Q67C+S59Y)-AMP in combination or not with pBAD-*dfrB1*-KAN. Then, from an overnight preculture grown in LB AMP (also + KAN for the duplication) medium, cells were diluted 1:100 in fresh medium with the addition of different amounts of arabinose, with concentrations ranging from 0 to 0.2%. Following an 18-hour incubation at 37°C (250 rpm), cells were diluted again at OD_600_ = 0.01 in a final volume of 200 μl of fresh medium containing arabinose with the addition of TMP or DMSO. The 96-well plates were incubated at 37°C in a BioTek Synergy HTX Multi-mode Reader (Agilent, USA) or in an Infinite M Nano plate reader (Tecan, Switzerland) for 13 to 16 hours. OD_600_ measurements were taken every 15 min. The plate was agitated at 237 cpm (4 mm) at slow orbital speed (Agilent) or at 200 rpm (Tecan) in between measurements.

The area under each growth curve was calculated using Growthcurver^73^. Growth recovery at each arabinose concentration with TMP relative to the WT in the absence of TMP was calculated using the following equation:

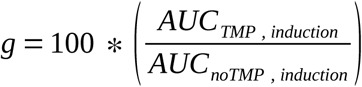

where *g* refers to the recovered growth percentage and *AUC* is the area under the curve in the conditions specified by the subscript (with or without TMP and with a given concentration of arabinose).

We used ANOVAs to analyze the differences in growth recovery and tetramer stability across mutants. In each case, the ANOVA was followed by Tukey’s honestly significant difference post hoc test computed using the agricolae R package^74^.

## Supplementary figures

**Figure S1:**
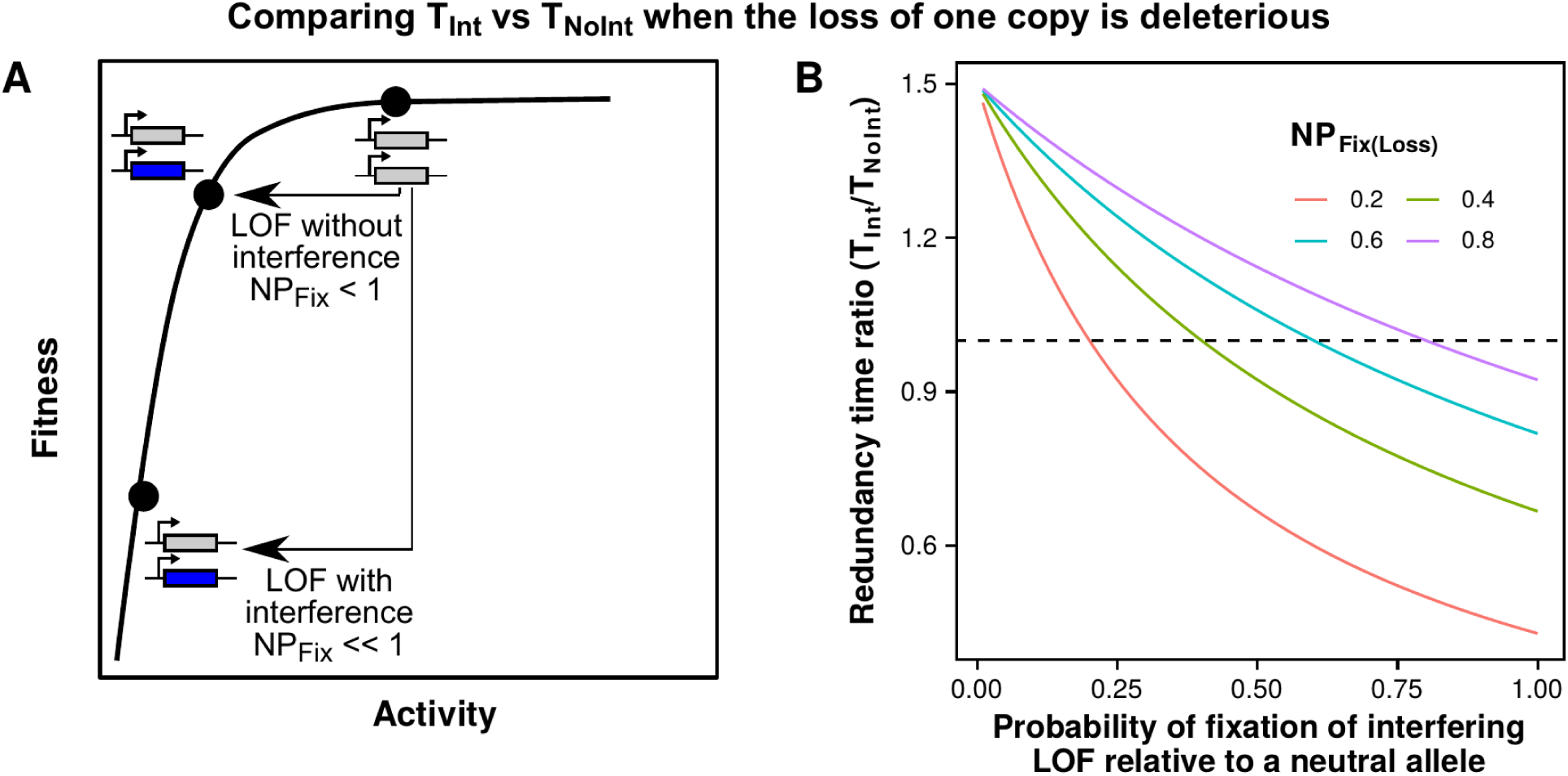
The increase in residence time of redundant paralogs due to interference decreases when a fraction of protein copies remain as monomers. A) Fitness function with diminishing fitness returns for increases in activity. A reduction of 50% in activity (LOE, LOA, and LOF mutations without interference) is deleterious, while stronger decreases (LOF mutations with interference) are more deleterious. B) Ratio of the time needed to lose redundancy in a system with interference (*T_Int_*) versus a system without interference (*T_NoInt_*). The ratio of mutation rates (*µ_A_*+*µ_E_*)/*µ_F_* is kept constant at 2. Different curves indicate varying magnitudes of deleterious effects for the loss of one copy. As shown in the Methods, T_Int_ / T_NoInt_ > 1 when the probability of fixation of a LOF mutation is lower than that of the loss of one copy.

**Figure S2:**
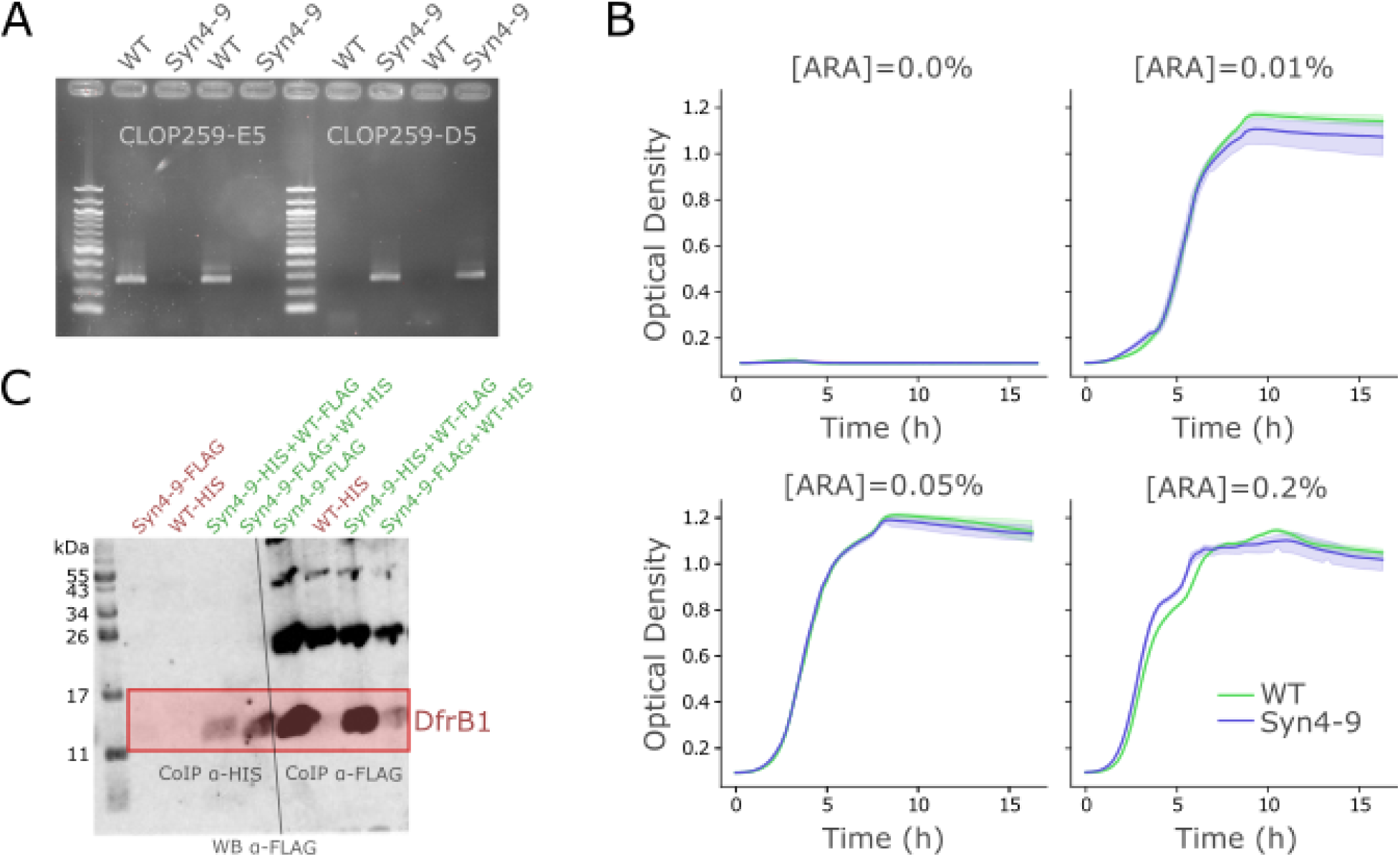
Synonymous mutations in the coding sequence of DfrB1 allows for specific amplification without impacting bacterial growth nor protein complex assembly. **A)** PCR validation of the differential amplification of WT *dfrB1* and the Syn4-9 variant, which allows distinguishing them during library preparation for deep sequencing. **B)** Growth curves showing there are no differences in the growth of WT and the Syn4-9 variant. **C)** Western blot after Co-IP to show the formation of DfrB1 heteromers when two gene copies are expressed, providing a system to study paralog interference. *E. coli* expressing the WT and the Syn4-9 mutant tagged with epitopes were used for these experiments. Samples were co-immunoprecipitated with ɑ-HIS or ɑ-FLAG as indicated in the two groups and then immunodetected for the presence of ɑ-FLAG. Samples are labeled at the top in green if DfrB1 is expected to be observed and in red otherwise. Bands observed in the Co-IP with ɑ-HIS indicate the presence of heteromers.

**Figure S3:**
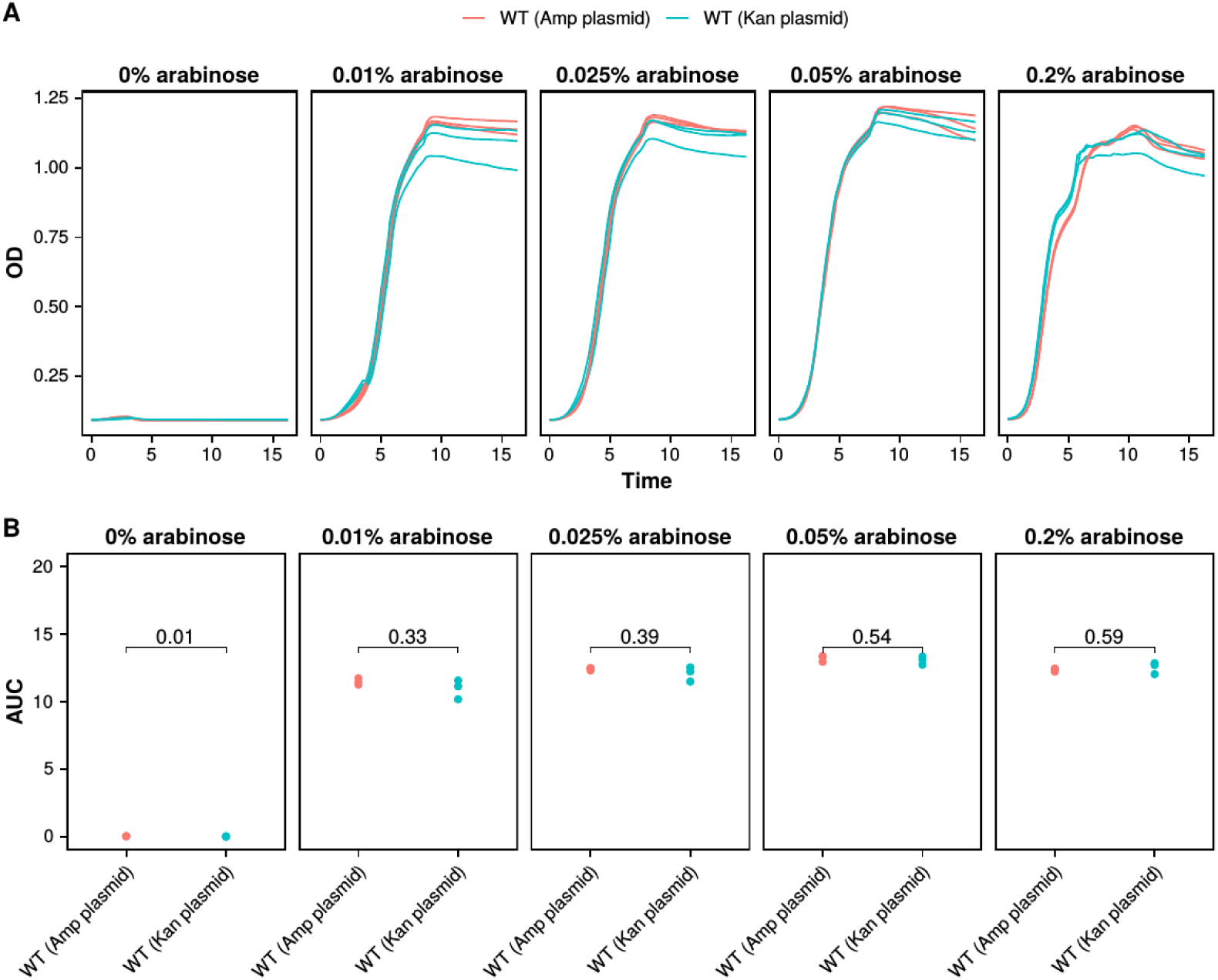
Expressing WT DfrB1 from two different plasmids results in similar growth. A) Growth curves for WT DfrB1 expressed from a plasmid harboring the Amp resistance marker vs one harboring the Kan resistance marker. B) Comparison of the area under the curve for each plasmid and arabinose concentration. p-values were obtained using a t-test.

**Figure S4:**
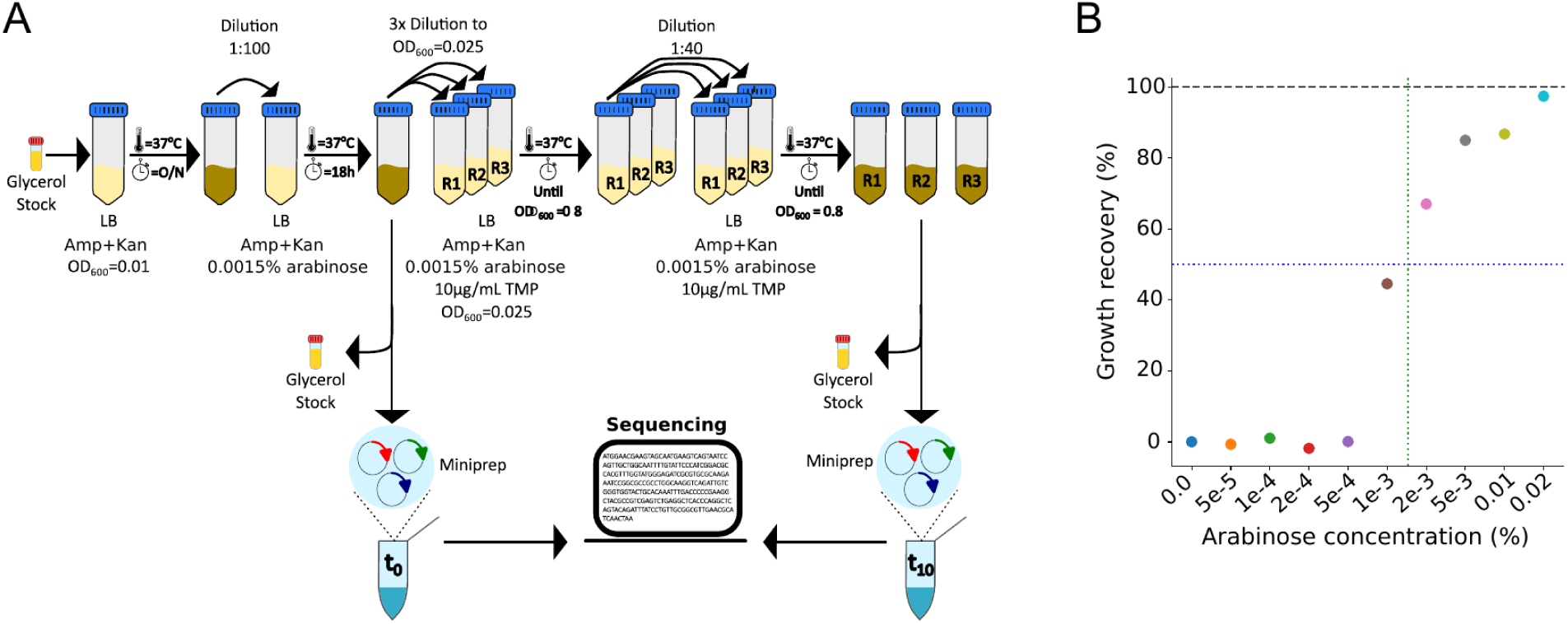
Selection of induction level (arabinose concentration) and design of the screening experiments using the mutant libraries. **A)** Overview of the bulk competition experiments. A preculture with the desired population is allowed to grow for 18 hours. It is then diluted to an optical density (OD) of 0.025 in three replicate populations, with an aliquot preserved for sequencing to estimate variant frequencies (t = 0). The three replicates are then allowed to grow at 37°C until they reach an OD = 0.8 (five generations). Each replicate is then diluted 1:40 and allowed to grow for five more generations. At t = 10 generations, an aliquot from each replicate is taken for sequencing. Selection coefficients are inferred based on the read counts for each mutation at t = 0 and t = 10, normalized by the read counts for the WT. **B)** The induction of DfrB1 expression with arabinose in media with TMP restores *E. coli* growth. Growth recovery is quantified as the area under the curve (AUC) at a given concentration of arabinose and 10 µg/mL TMP, normalized by the AUC without TMP.

**Figure S5:**
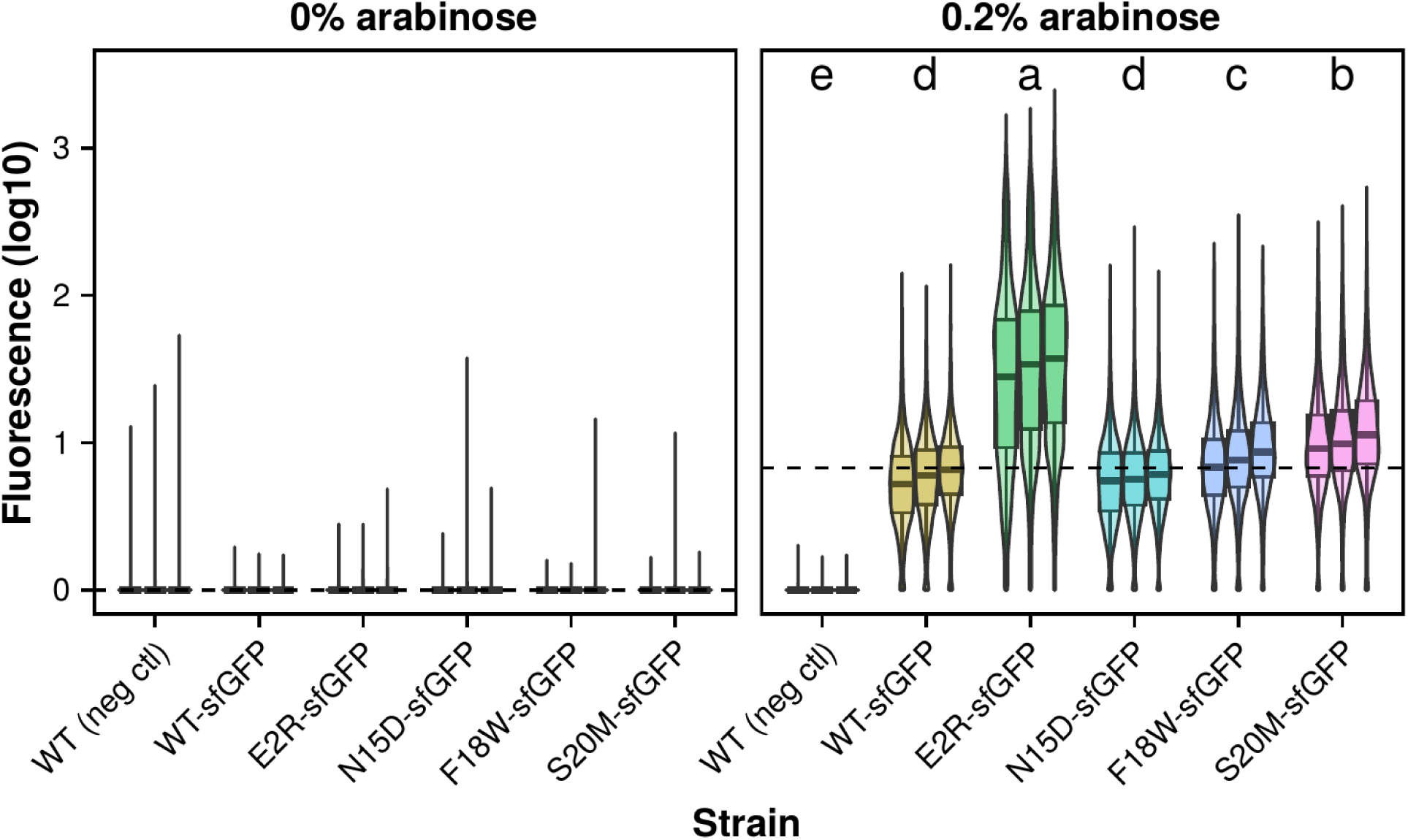
Some substitutions in the disordered region of DfrB1 that increase fitness do so by increasing protein abundance. Substitutions in the disordered region with beneficial fitness effects (E2R, N15D, F18W, S20M) were selected and fused to sfGFP. Comparison of fluorescence values observed in flow cytometry for three replicate populations of ∼5000 cells harboring each mutant. Dashed lines indicate the median of the WT-sfGFP fluorescence for the pool of cells from the three replicates. Labels at the top indicate groups of fluorescence shown to be significantly different using an analysis of variance (ANOVA) and Tukey’s post hoc honestly significant difference test (p < 0.05).

**Figure S6:**
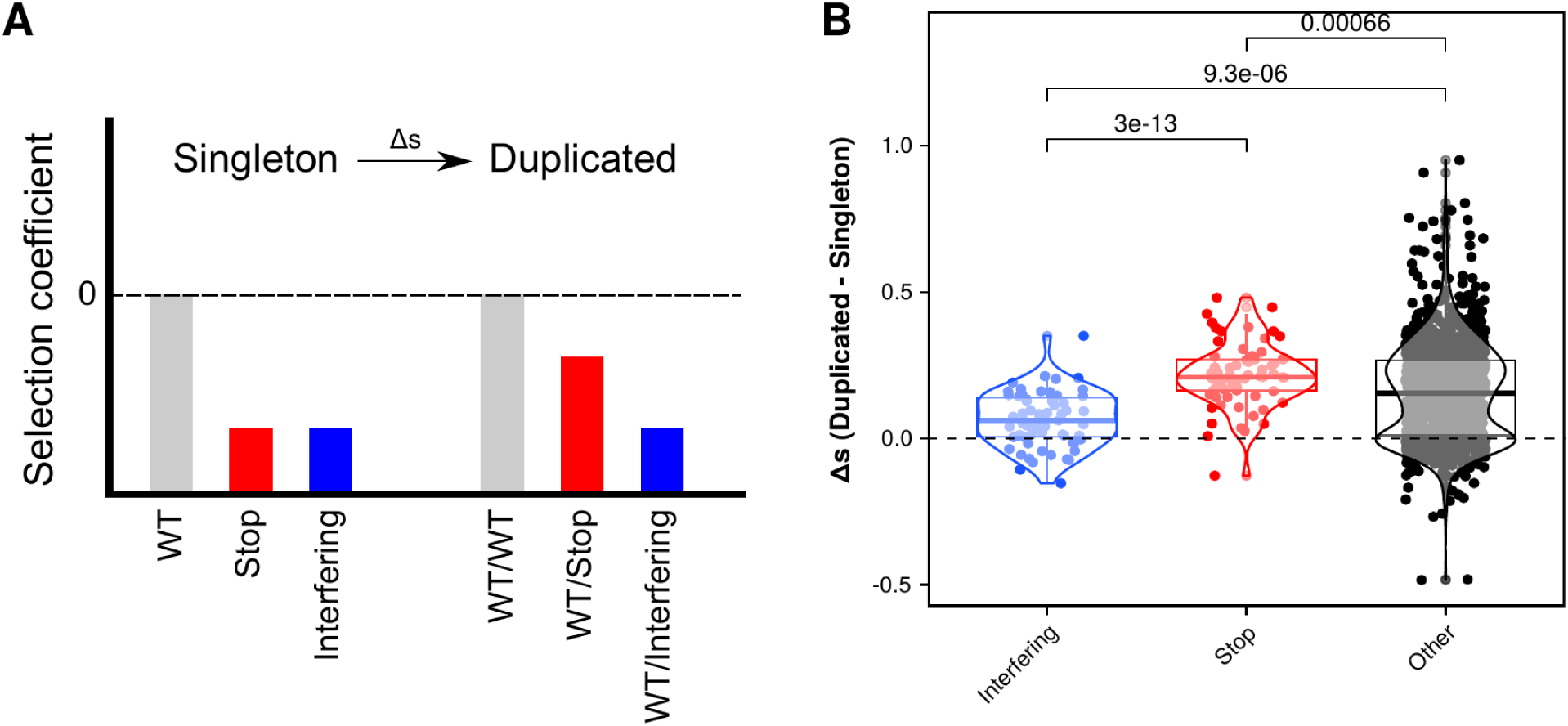
The deleterious effects of interfering substitutions are not rescued by a second WT copy. A) Expectations of selection coefficients for stop codons and interfering substitutions in the singleton and background scenarios. In the singleton scenario both types of substitutions are equally deleterious, while in the duplicated scenario interfering substitutions are more deleterious than stop codons. B) Differences in selection coefficients observed for different types of substitutions in the duplicated versus singleton scenario. Positive values indicate those that are less deleterious in the duplicated scenario than in singleton. p-values were calculated using Wilcoxon tests.

**Figure S7:**
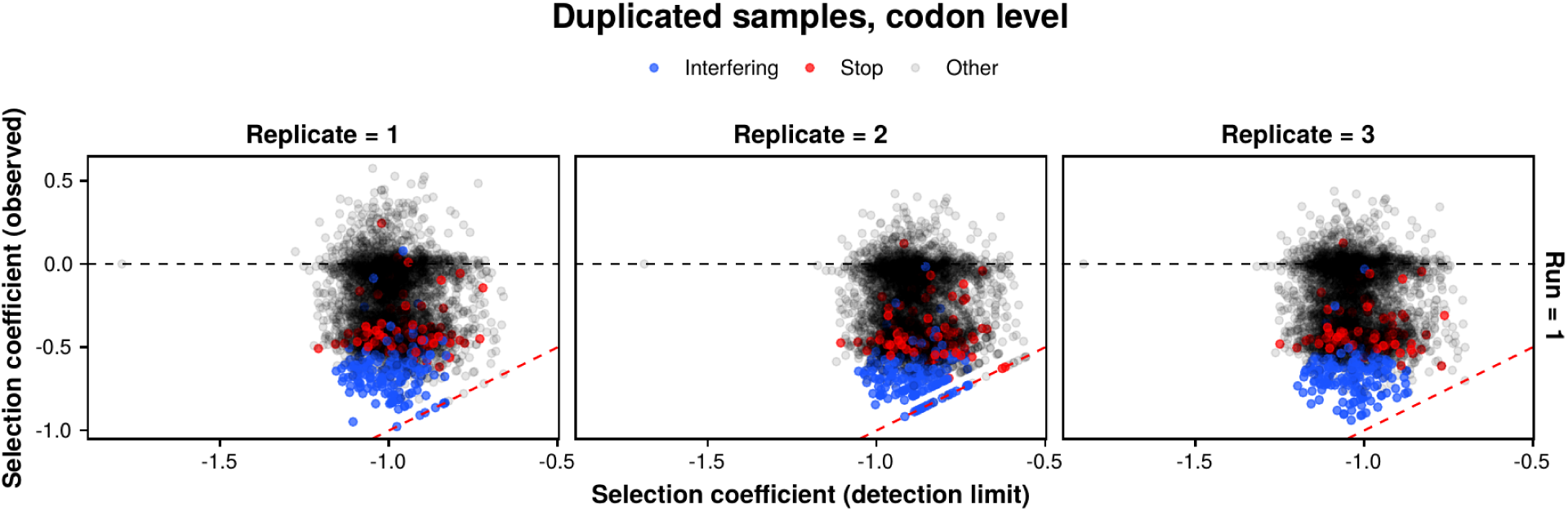
Observed selection coefficients are within the dynamic range provided by pool composition. Observed selection coefficients (y-axis) are compared versus the most deleterious selection coefficient observable for each mutant (x-axis). The most deleterious selection coefficient observable corresponds to the mutant going from its initial read frequency to zero at the end of the experiment. The horizontal dashed line indicates the selection coefficient for the WT (s = 0), while points on the red dashed line indicate substitutions whose observed selection coefficients match the most deleterious observable values. The three replicates of the experiment with the duplicated background are shown separately.

**Figure S8:**
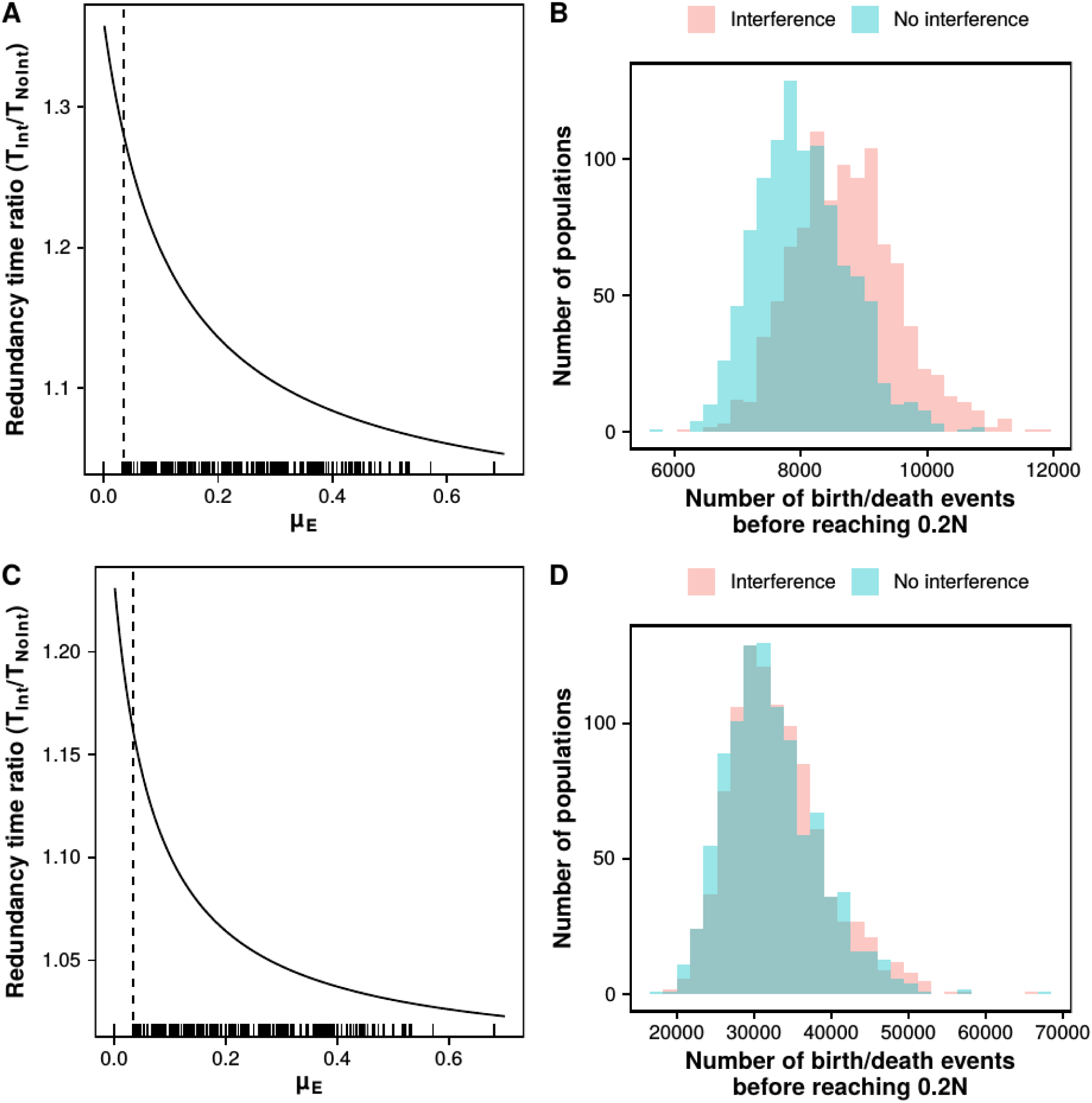
Increase in residence time of redundant paralogs with similar properties to DfrB1. A-D) Ratio of residence times of redundant paralogs with mutation rates similar to DfrB1 with and without interference was estimated analytically (A, C) and using a Moran model (B,D). **A, C)** For analytical estimates, the curve was calculated using the proportions of interfering (LOF-like) and mutations destabilizing the dimerization interface (LOA) with varying rates of LOE mutations. All sampled mutations were considered for the estimation of mutation rates in panel A, whereas only those accessible by a single point mutation were considered in panel C. Rugs under panels A and C indicate the rates of LOE mutations observed for 2034 promoters by Urtecho et al.^31^, with the vertical dashed line indicating the median value of 0.034. **B, D)** Number of birth/death events required in a Moran model to reduce the proportion of individuals harboring two redundant copies of paralogs to 0.8N. Simulations were carried out with and without interference using the same parameters from panel A for DfrB1 and the rates of LOE mutations from Urtecho et al.^31^ for a sample of 1000 populations with N = 1000 individuals each. The same parameters as in panels A and C were respectively used for panels B and D.

**Figure S9:**
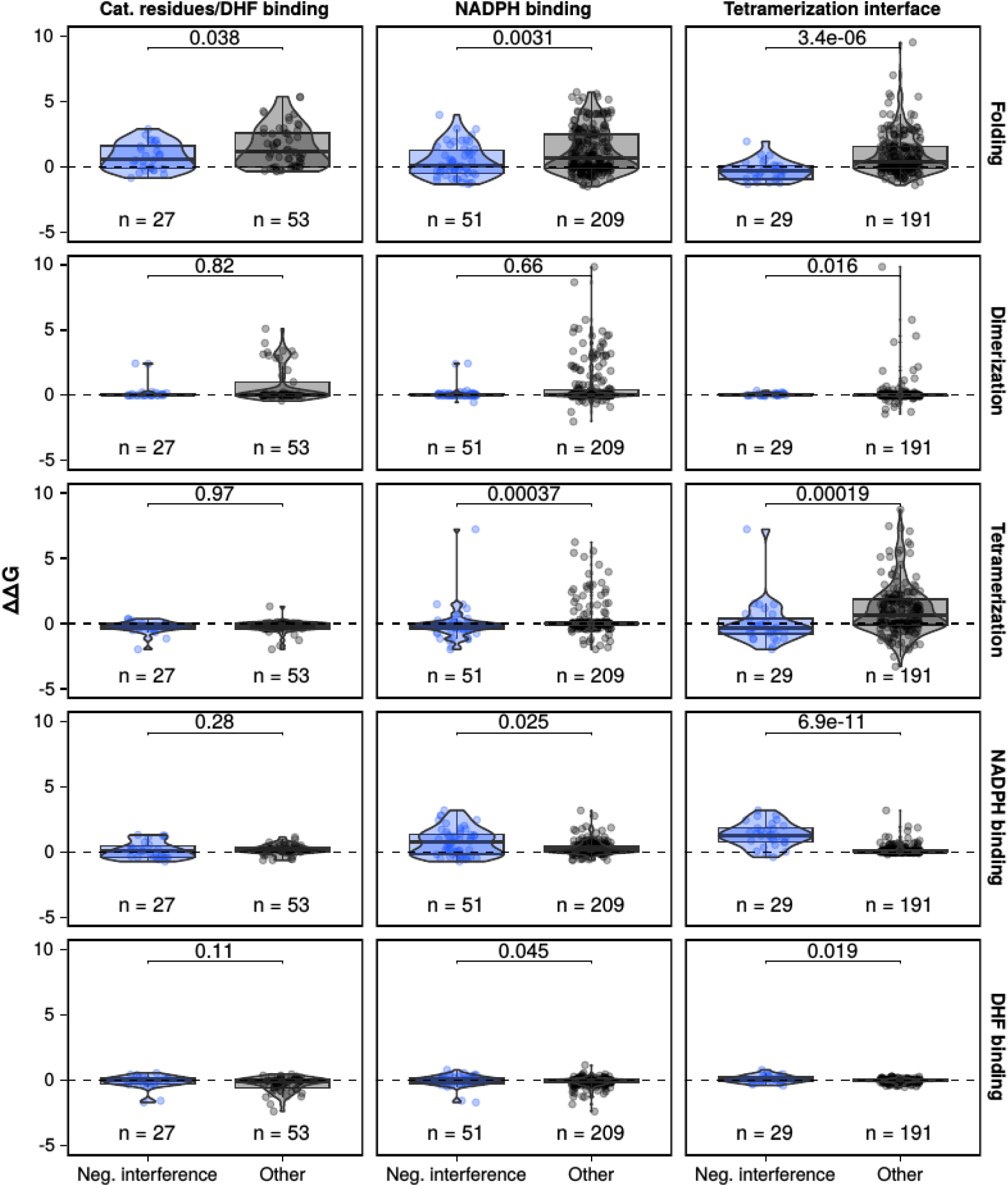
Negative interference substitutions in specific protein regions have milder effects on stability and stronger effects on NADPH binding than other substitutions in the same region. Same data as in Figure 3B, but filtered to show only substitutions occurring in the regions with the highest enrichment of negative interference substitutions in Figure 3C (columns). Rows indicate the different biophysical effects. p-values were calculated using Mann-Whitney U tests. As shown in Fig. 3C and^24^, some positions are assigned to more than one region. Effects on NADPH binding of substitutions in the tetramerization interface could be associated with positions that are directly in contact with NADPH or adjacent to residues that are in contact with NADPH.

**Figure S10:**
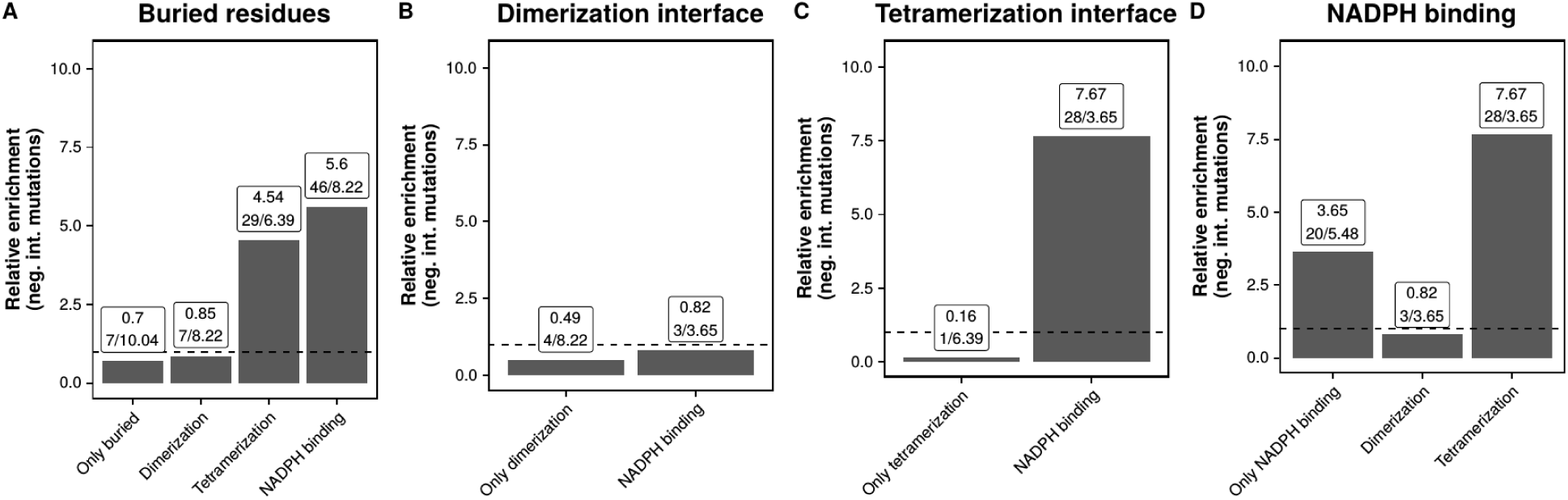
Relative enrichment of interfering substitutions across selected deconvoluted regions. A-D) Using the same data as in Figure 3C, the relative enrichment of interfering substitutions was recalculated with emphasis on buried residues (A), the dimerization interface (B), the tetramerization interface (C), and the NADPH binding sites (D). In each panel, the residues belonging to the corresponding region were separated as those belonging to only that region and those overlapping with each of the other regions. The dashed line indicates a relative enrichment of 1, in which the observed counts of negative interference substitutions are identical to the expected counts. Numbers above bars indicate the observed relative enrichment of interfering substitutions, calculated as a ratio of observed over expected counts.

**Figure S11:**
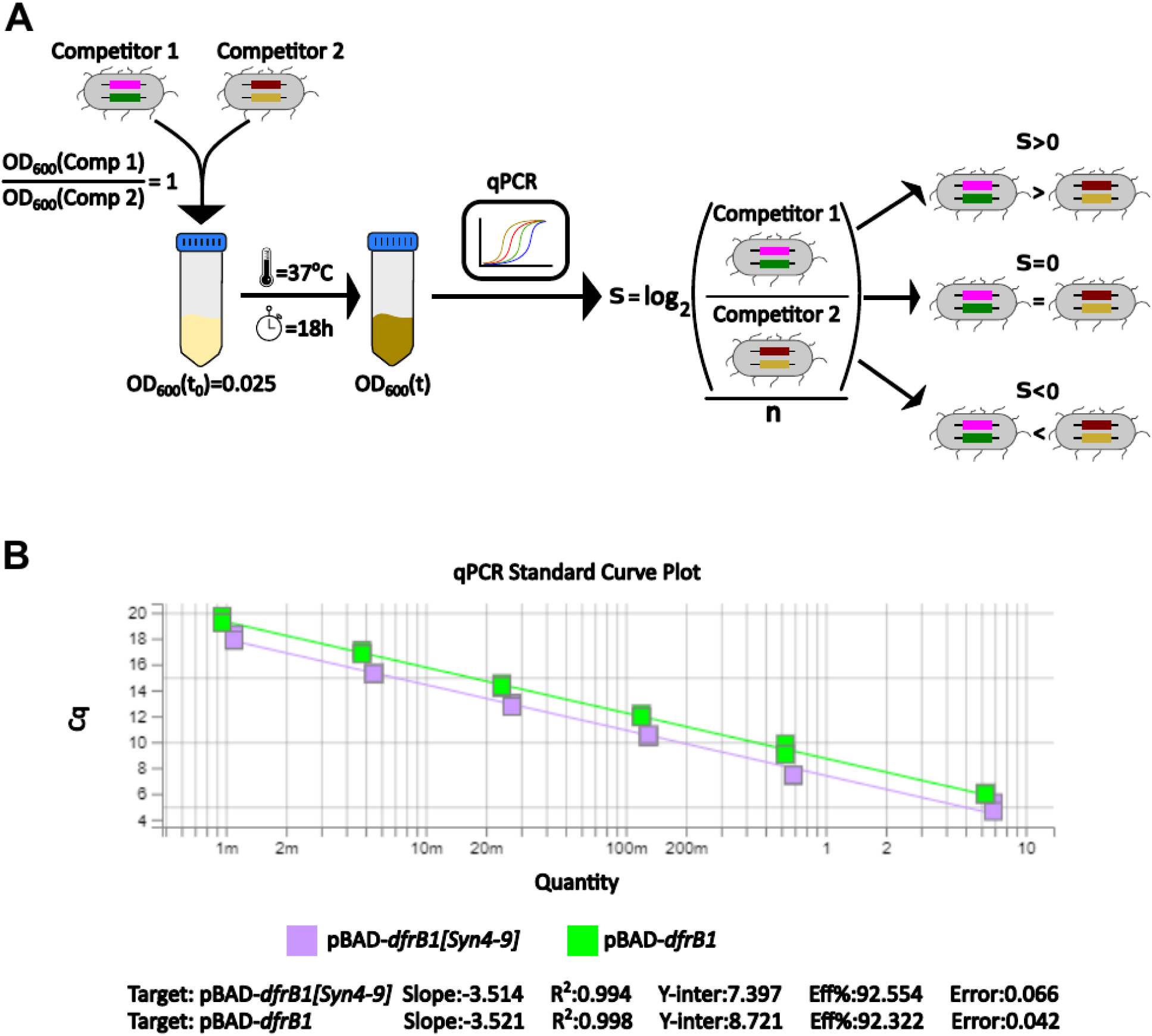
Direct competition experiment and measurement of relative abundance using qPCR. A) In each experiment the two competitors, one with the Syn4-9 mutation and the other without, were inoculated at the same proportion based on the individual OD_600_ and grown together for 18h. The number of generations (n=log_2_[OD_600_(t)/OD_600_(t0)]) was determined by measuring the OD_600_ of the grown culture. 1 μL was directly used to quantify selection coefficients (s) of one competitor relative to the other targeting the change in proportions for the WT and the Syn4-9 sequence by quantitative PCR. B) Standard curve showing the relationship between amplification cycles (Cq) values and log-transformed template concentrations. Both targets showed linearity and similar efficiency (∼92%), confirming reliable quantification for all the samples tested across the experiment.

**Figure S12:**
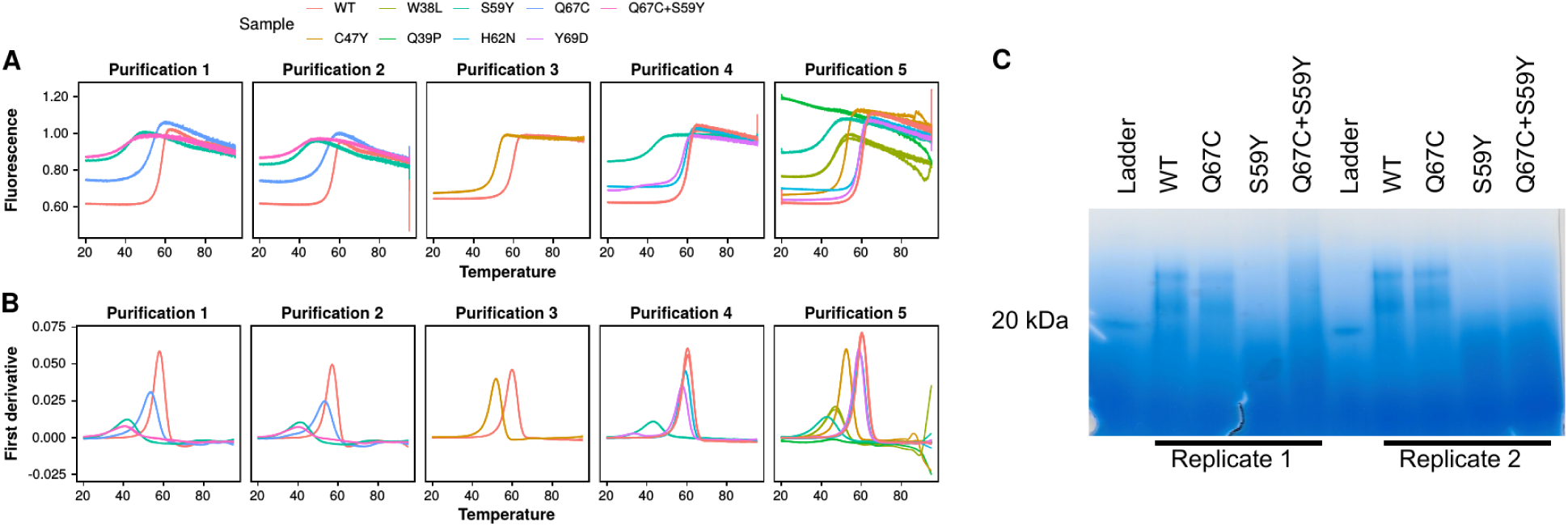
Effects of selected substitutions on DfrB1 assembly. A-B) Denaturation curves shown as the ratio of emitted fluorescence at 350 nm vs 330 nm as a signal of exposed tryptophan residues (A) and their first derivatives (B) for different DfrB1 variants. All denaturation curves were obtained with three replicates for each variant. Data were obtained across five separate purification experiments comprising different lists of variants. **C)** Native gel electrophoresis of purified DfrB1 mutants. Labels indicate samples loaded in the corresponding wells.

**Figure S13:**
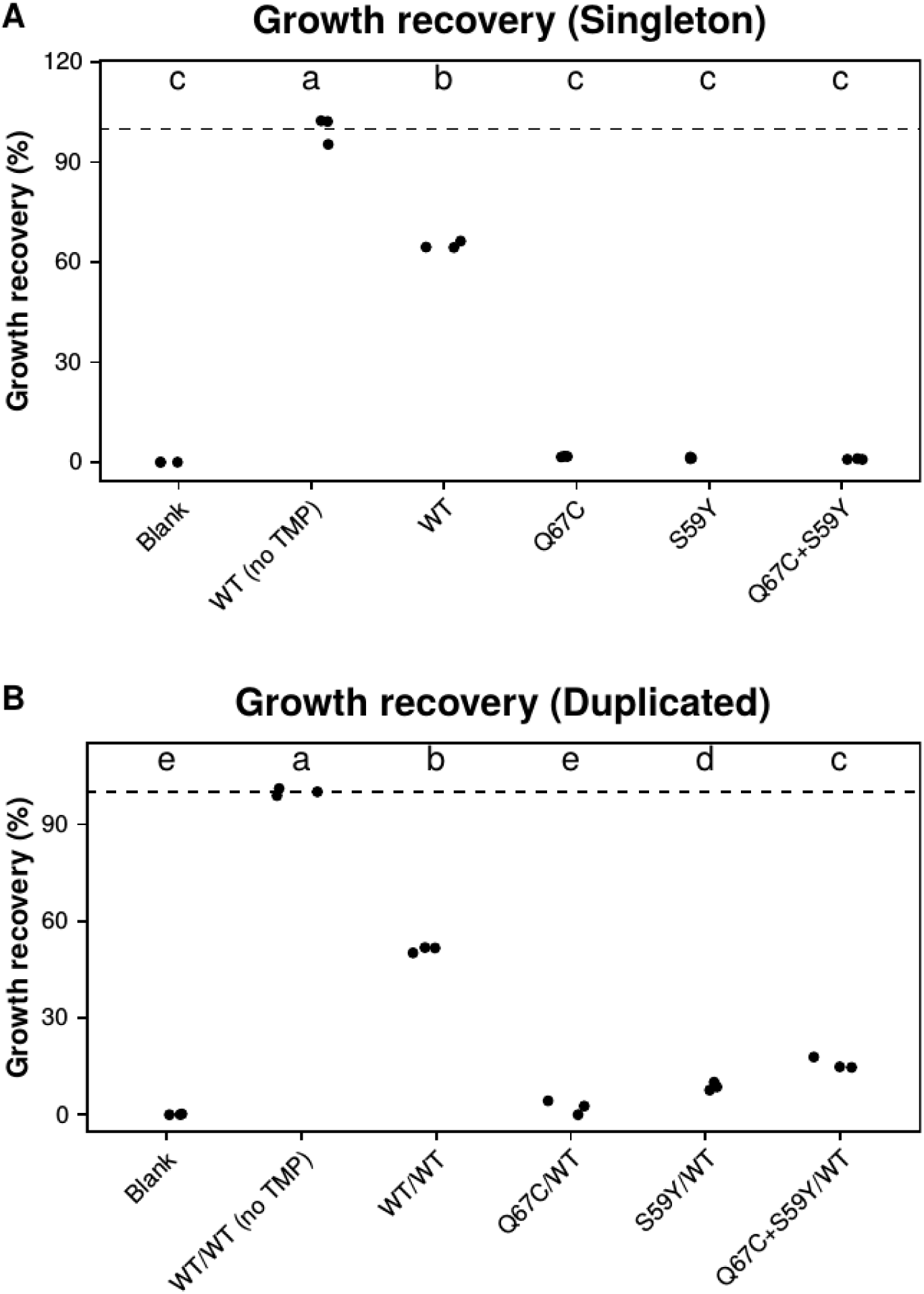
Selected negative interference substitutions cannot restore growth either as a singleton or alongside a WT copy. A-B) Comparison of growth recovery at 0.0015 % arabinose and 10 µg/mL TMP for different genotypes in single copy relative to a WT reference growing in the absence of TMP (A) and alongside a WT copy relative to a WT/WT reference growing in the absence of TMP. Labels at the top indicate groups of recovered growth shown to be significantly different using an analysis of variance (ANOVA) and Tukey’s post hoc honestly significant difference test (p < 0.05).

